# Targeted Deletion of All Known Thyroid Hormone Receptors Causes Maturation Retardation and Early-onset Degeneration of Cochlear Outer Sulcus

**DOI:** 10.1101/2023.10.31.565054

**Authors:** XianHua Ma, Fei Jiang, Chunchun Wei, Shuang Han, Yuqing Zhang, Lianhua Sun, Jiaxi Qu, Hao Ying, Yuxia Chen, Jie Tang, David Z. He, Weiping J. Zhang, Zhifang Xie

## Abstract

Thyroid hormone (TH) and its receptors (TRs) are crucial for cochlear late-stage development and endocochlear potential (EP) maintenance. However, the mechanism underlying EP reduction in the absence of TH or TRs remains elusive. Cochlear outer sulcus root cells undergo significant morphological changes during late-stage cochlear development and are believed to play a role in maintaining endolymph homeostasis and EP. Yet, it is unknown whether TH and/or TRs are necessary for root cell differentiation and function. Here, we elucidate the essential role of TH or TRs in postnatal root cell development and survival in mice. TH deficiency significantly delayed root cell differentiation. Otocyst-selective deletion of both *Thra* and *Thrb*, but not *Thrb* alone, leads to a similar impairment, accompanied by early degeneration of root cells, with the stria vascularis remaining unaffected. Furthermore, a 22% reduction in mean EP magnitudes was observed in conditional *TRs* double knockout mice at 4 months of age, less pronounced than in global *TRs* knockout mice. Transcriptome analysis reveals that TH deficiency downregulates a significant portion of root cell-enriched genes. These findings underscore the redundant roles of TRα and TRβ in promoting the late-stage differentiation and survival of root cells. Additionally, they suggest that the expression of TRs in cochlear epithelium is crucial for maintaining an optimal EP magnitude, while TRs expressed in areas outside cochlear epithelium, particularly in spiral ligament fibrocytes, may also significantly contribute to EP maintenance. This study advances our understanding of TH in cochlear outer sulcus development and EP maintenance.

## Introduction

Thyroid hormone (TH) is crucial for the normal development of the cochlea and hearing (Ng et al., 2013). Both early-onset congenital hypothyroidism and environmentally induced iodine deficiency can lead to hearing loss in both humans and rodents (Christ et al., 2004; Deol, 1973a; Mustapha et al., 2009). TH primarily exerts its effects by binding to two distinct thyroid hormone receptors (TRs), namely TRα and TRβ, which are ligand-dependent transcription factors encoded by *Thra* and *Thrb*, respectively (Forrest et al., 1996; Rusch et al., 2001; Winter et al., 2009). TRα is individually dispensable for hearing(Rüsch et al., 1998), while the absence of TRβ alone leads to deafness accompanied by malformation of the tectorial membrane (TM) (Forrest et al., 1996; Winter et al., 2009). Global deletion of both *Thra* and *Thrb* in mice exacerbates cochlear defects, including more severe distortion of the TM, significant retardation in the regression of the great epithelial ridge (GER, also known as Kolliker’s organ), and remodeling of the organ of Corti, as well as a reduction in endocochlear potential (EP) (Rusch et al., 2001). The mechanism underlying the reduction in EP upon deletion of TRs remains unclear, as no significant alterations in the stria vascularis (SV) have been reported in these mice, suggesting that defects beyond the SV may contribute to the reduction in EP.

The cochlear outer sulcus epithelial cells, also known as root cells, are poorly understood cell types that are considered to be important for the maintenance of endolymph homeostasis and EP (Spicer and Schulte, 1996; Cazals et al., 2015; Galic and Giebel, 1989; Jagger et al., 2010; Jagger and Forge, 2013). Positioned on the cochlear lateral wall between the outer border Claudius cells and the spiral prominence, the outer sulcus undergoes significant morphological changes during the postnatal developmental window when the cochlea is most sensitive to TH (Deol, 1973b; Karolyi et al., 2007; Uziel et al., 1985). At birth, the outer sulcus consists of a simple layer of cuboidal-shaped epithelial cells lining the scala media. Shortly after birth, these cells begin lateral and superior infiltration into the spiral ligament beneath the spiral prominence, forming extensive tree-like branched processes, hence termed “root cells.” Concurrently, Claudius cells enlarge significantly, meeting the spiral prominence epithelium and covering the underlying root cell bodies, thereby preventing root cells from direct contact with endolymph in basal turns. However, in the apical turn, root cells directly face the endolymph and exhibit fewer processes (Duvall III, 1969; Galic and Giebel, 1989; Jagger and Forge, 2013; Lim and Anniko, 1985; Shodo et al., 2017; Spicer and Schulte, 1996).

There is limited understanding of the mechanisms that regulate root cell differentiation. Our recent study has highlighted the importance of the transcription factor ZBTB20 in cochlear late-stage development, particularly in root cell differentiation. Selective deletion of *Zbtb20* in the cochlear epithelium results in deafness reduced EP, and cochlear malformations, such as a malformed tectorial membrane (TM) and organ of Corti, along with impaired maturation of the outer border and outer sulcus region (Xie et al., 2023). Intriguingly, *Thrb* expression has been detected in both developing and adult root cells, as demonstrated by a lacz gene reporter (Ng et al., 2015) or single-cell RNA sequencing (Gu et al., 2020), while *Thra* mRNA is widely expressed in the lateral wall, including root cells (Bradley et al., 1994; Jean et al., 2023). However, it has long been neglected whether TH and the expression of TRs in the cochlea play essential roles in the remodeling and maturation of the outer sulcus.

Here we find that TRs expression in otocyst-derived cochlear epithelium is necessary for postnatal root cell development and survival. Root cell development is significantly delayed in mice with antithyroid drug-induced developmental hypothyroidism or those with otocyst-selective deletion of both *Thra* and *Thrb*, while no apparent root cell defects are observed upon individual deletion of *Thrb*. Moreover, the absence of both TRα and TRβ results in early-onset degeneration of root cells and a compromised EP magnitude.

## Results

### Hypothyroidism causes severe retardation in the outer sulcus development

To investigate the role of thyroid hormone (TH) in outer sulcus development, we induced developmental hypothyroidism through antithyroid treatment, following a previously established protocol(Sawant et al., 2015). Antithyroid treatment commenced on gestation day 17 and extended through lactation until postnatal day (P) 21. This duration covers a critical developmental window during which the cochlea exhibits heightened sensitivity to TH deprivation (Deol, 1973b; Uziel et al., 1985). Histological analysis was conducted on cochleae harvested from euthyroid or hypothyroid mice at P21, a stage when the cochlea attains both morphological and functional maturity in normal mice (Lim and Anniko, 1985; Walters and Zuo, 2013).

To better preserve cochlear histological features, we employed a resin-embedding technique. In euthyroid cochleae, the sensory epithelium exhibited typical mature morphology, characterized by an upright organ of Corti, a well-developed inner sulcus, and a normal tectorial membrane (TM). Within the outer sulcus region, enlarged Claudius cells covered the underlying root cells, displaying numerous bright branched processes extending deep into the spiral ligament and upwards behind the spiral prominence (Fig. 1A). In line with previous findings(Deol, 1973b), hypothyroid cochlear epithelium exhibited marked retardation in sensory epithelium remodeling, evidenced by the abnormal persistence of the greater epithelial ridge (GER), closure of the tunnel of Corti, and severe distortion of the TM. Notably, the outer sulcus displayed an immature appearance, characterized by a simple layer of cells with few short processes bordering the scala media, along with flattened Claudius cells (Fig. 1B). This observation suggests severe retardation in both root cell and Claudius cell differentiation.

**Fig. 1.**
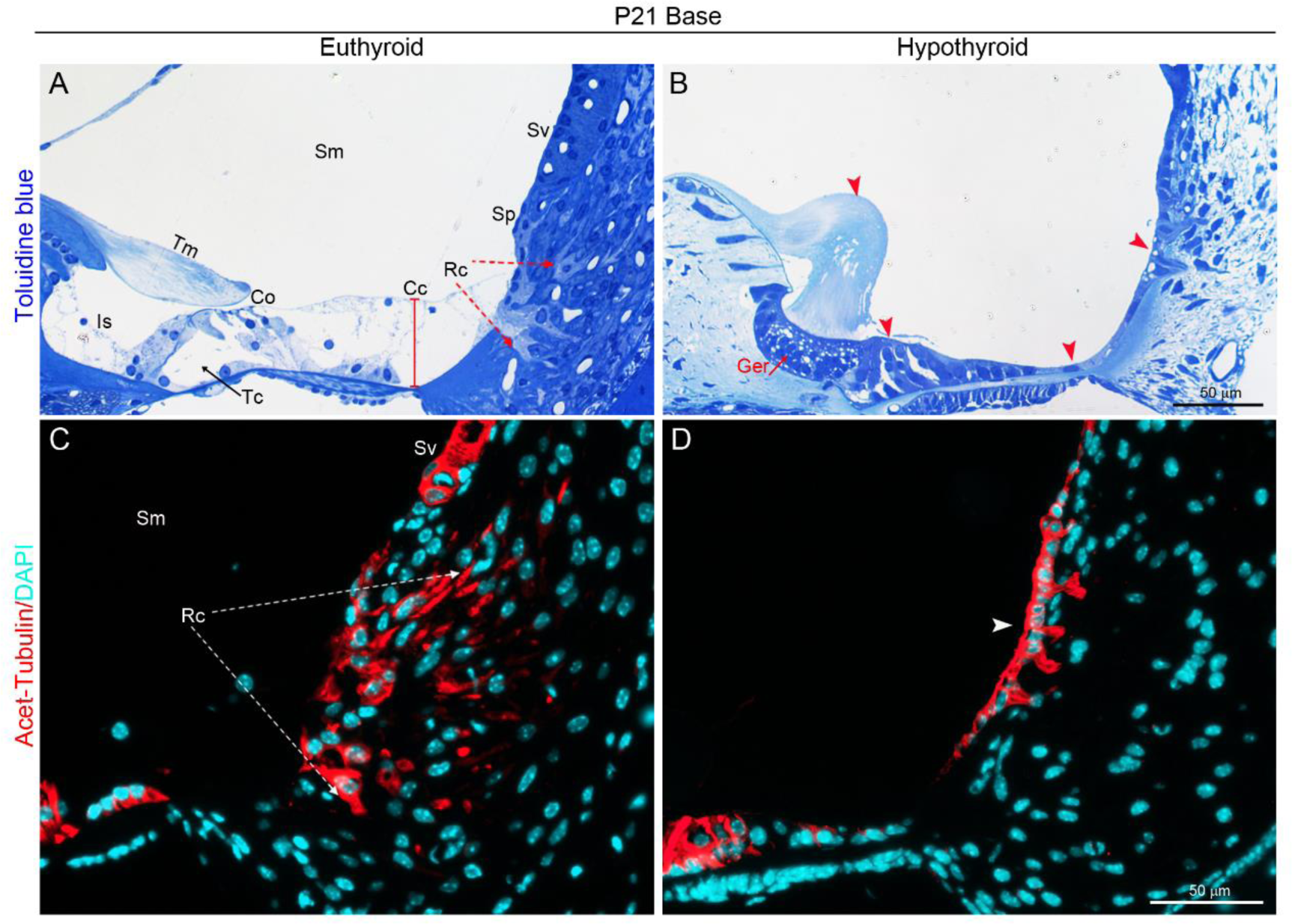
Hypothyroidism disrupts the outer sulcus development. **(A and B)** Representative Toluidine Blue-stained sections of cochlear upper basal turn from euthyroid (**A**) or methimazole (MMI)-induced hypothyroid mice (**B**) at P21. (**A**) Euthyroid cochlea showing a mature outer sulcus characterized by root cells (Rc. indicated by red arrows) with numerous bright branched processes extending deep into the spiral ligament and upward to behind the spiral prominence (Sp). The surfaces of root cell bodies were covered by enlarged Claudius cells (Cc, red bar indicating its height). Co, the organ of Corti; Is, inner sulcus; Sm, scala media; Sv, stria vascularis; Tc, tunnel of Corti, Tm, tectorial membrane. (**B**) The hypothyroid outer sulcus consists of a simple layer of cells with a few short processes bordering the scala media, continued with the flattened Claudius cells. Red arrowheads indicate defects including a distorted Tm, an abnormal persistence of GER, closure of tunnel of Corti, and a flattened border region. (**C** and **D**) Immunostaining shows extensive acetyl-α-tubulin positive (red) root cell processes extending into the inferior region of the euthyroid spiral ligament at P21 (**C**) but only a few short processes observed in the luminal surface of hypothyroid counterparts (indicated by white arrowhead) (**D**). Bars = 50 µm for all panels. Nuclei were stained with DAPI (turquoise). n = 4 mice/ group. Scale bars in all panels = 50 µm. Abbreviations are consistent across figures in this manuscript.

Our previous work, along with studies by others, has demonstrated that root cell cytoplasm and their processes exhibit enrichment of acetyl-α-tubulin (Jagger and Forge, 2013; Liu et al., 2018; Xie et al., 2023). To gain further insight into root cell morphology, we conducted immunofluorescence staining for acetyl-α-tubulin. Consistent with the histological findings, euthyroid cochleae exhibited robust and extensive acetyl-α-tubulin signals extending deep into the spiral ligament (Fig. 1C). In contrast, hypothyroid cochleae displayed limited signals, primarily distributed around the luminal surface of the spiral ligament, indicative of root cell differentiation defects (Fig. 1D). These findings underscore the essential role of TH signaling in the timely development of the outer sulcus.

### Otocyst-selective deletion of all known *TRs* causes deafness and a compromised EP

As TH effects are mainly mediated by TRα and TRβ(Ng et al., 2013), both of which are expressed in the developing outer sulcus(Bradley et al., 1994; Gu et al., 2020; Ng et al., 2015), we therefore asked if the two receptors are essential for root cell differentiation. To address this question, we generated mice with either individual deletion of the *Thrb* gene or combined deletion of the *Thra* and *Thrb* genes selectively in the otic vesicle using the Cre-loxP system. These mice are hereafter referred to as OV-TRβKO and OV-TRα/βKO, respectively. Individual *Thra*-deleted mice were not included in this study, as *Thra*-null mice have been reported to exhibit normal hearing thresholds (Rüsch et al., 1998). We crossed *Thrb-floxed* (*Thrb^flox^*) mice(Yan et al., 2022) either individually or in combination with *Thra ^flox^* mice(Liu et al., 2022) to *Foxg1-Cre* transgenic mice, a strain capable of mediating recombination selectively in the otic vesicle by embryonic day 8.5(Hebert and McConnell, 2000). We verified the tissue specificity of *Cre*-mediated recombination by mating *Foxg1*-*Cre* mice with *Rosa26*-lacZ reporter mice. Consistently, *Cre*-recombinase activity, as indicated by X-gal staining, was confined to otocyst-derived cochlear epithelium at P5. This included the greater epithelial ridge (GER), lesser epithelial ridge (LER), outer sulcus epithelium (root cells), stria vascularis (SV) marginal cells, and spiral ganglion. As anticipated, no signal was detected in cochlear fibrocytes derived from the periotic mesenchyme surrounding the otocyst (Song et al., 2011). No background signal was observed in cre-negative control mice (Fig. 2A). The efficiency of deletion was further evaluated through qRT-PCR analysis, which revealed that the expression levels of *Thrb* mRNA were significantly downregulated in OV-TRβKO cochleae at P10, while both *Thrb* and *Thra* mRNA were significantly downregulated in OV-TRα/βKO cochleae (Fig. 2B).

**Fig. 2.**
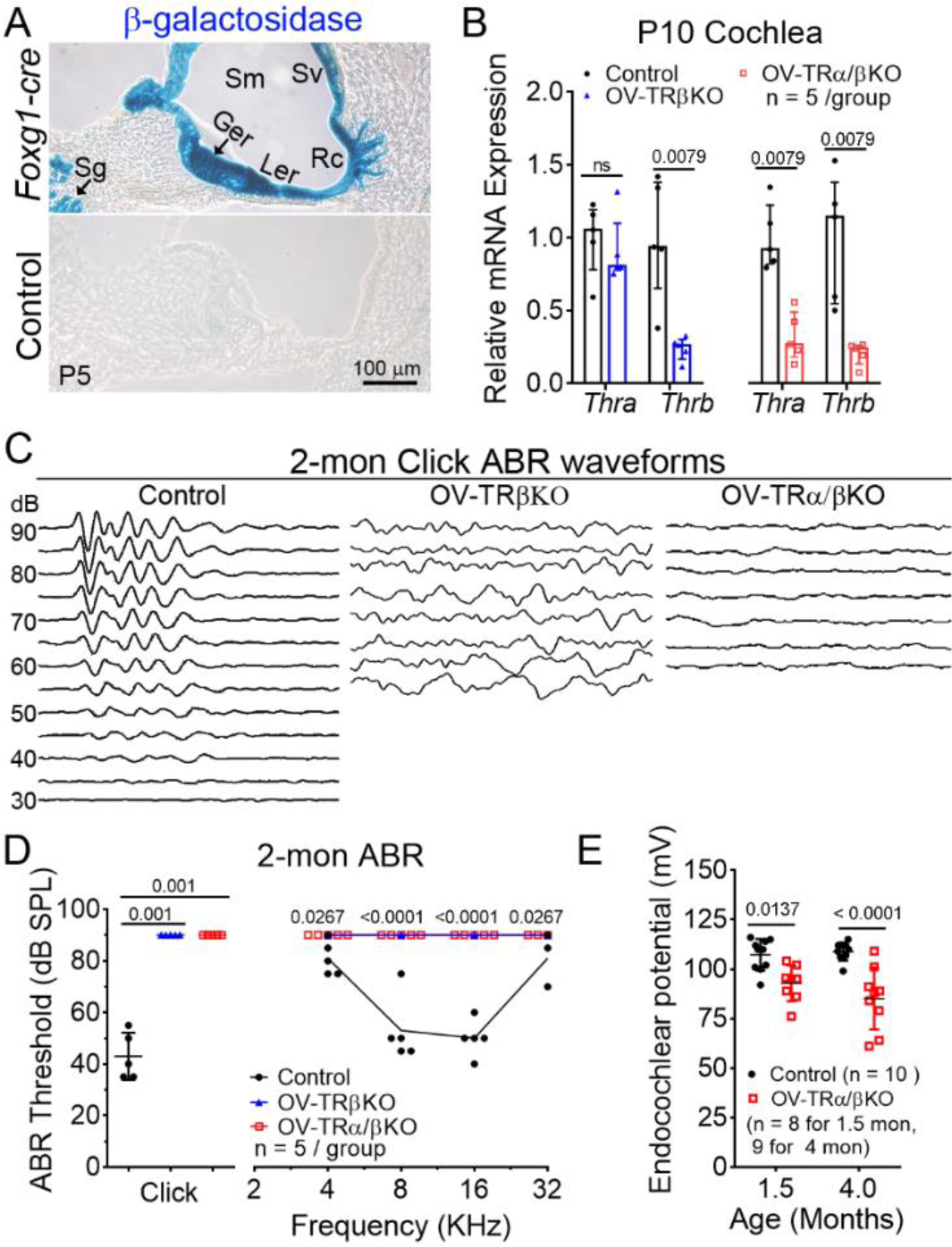
Otocyst-selective deletion of all known TRs causes deafness and a compromised EP. **(A)** X-gal staining showing *Foxg1-Cre* recombinase activity was confined in the cochlear epithelium lining the scala media (Sm) at P5, including GER, LER (the lesser epithelial ridge), root cells (Rc), stria vascularis (Sv), as well as spiral ganglions (Sg). n = 3 mice/group. Bar = 100 µm. **(B)** QRT-PCR analysis showing relative mRNA expression levels of *Thra* and *Thrb* in three groups of cochlea at P10. n = 5 mice/group. **(C)** Representative ABR waveforms obtained to broadband click stimuli from three groups of mice at 2 months (mon) of age. Sound pressure levels (SPL) were indicated in decibels (dB). (**D**) Averaged thresholds of Click- and Pure tone-evoked ABRs from three groups of mice at 2 mon. n = 5 mice/group. (**E**) OV-TRα/βKO mice showed a reduced EP magnitude in the basal turns compared with control mice at around 1.5 or 4 mon. The number of ears tested was indicated in the graph. Values are shown as medians with 25-75% interquartile ranges for (**B**), mean ± SD for (**D**), and (**E**). A Mann-Whitney test and an unpaired Student’s t-test were used to compare differences in (**B**) and (**E**), respectively. A Kruskal-Wallis test with Dunn’s multiple comparisons test and a two-way ANOVA followed by post-hoc pairwise tests (Bonferroni method) were used to compare differences in Click- and Pure tone-evoked ABR thresholds, respectively. *P* Values were indicated. ns, not significant.

To evaluate the auditory function of the mutant mice, we performed ABR (auditory brainstem response) tests. Both OV-TRβKO and OV-TRα/βKO mice did not show typical ABR responses to either click (Fig. 2C, D) or frequency-specific tone burst stimuli at 90 dB (the highest intensity tested) around 2 months (Fig. 2D), in sharp contrast to normal thresholds seen in wild-type (WT) control mice. These data demonstrate that cochlear expression of TRβ or all known TRs is essential for hearing, in agreement with previous reports(Forrest et al., 1996; Rusch et al., 2001).

The global deletion of all known *TRs* has previously been associated with a roughly 50% reduction in the EP magnitude in adult mice (Rusch et al., 2001). To investigate if the expression of TRα and TRβ within the cochlear epithelium is essential for EP maintenance, we measured EP in OV-TRα/βKO mice. At around 1.5 months of age, OV-TRα/βKO mice exhibited a slight but statistically significant decrease in EP magnitude compared to WT controls (WT = 107.2 ± 8.0 mV vs OV-TRα/βKO = 92.8 ± 9.0 mV, p = 0.01). By 4 months of age, the mean EP magnitude in OV-TRα/βKO mice further declined to 85.1 ± 15.6 mV, while no significant changes were observed in the WT group (108.8 ± 4.6 mV) (Fig. 2E). These findings underscore the role of TRs expressed within the cochlear epithelium in maintaining optimal EP magnitude. Additionally, they suggest that TRs expression in areas outside of the cochlear epithelium may also contribute to EP maintenance.

### Otocyst-selective deletion of all *TRs* retards root cell development

We next investigated cochlear morphological changes in *TRs* knockout mice, with an emphasis on the development of the outer sulcus region. At P10, 2 days before the onset of hearing in mice, root cells with branched processes had already invaded deep into the spiral ligaments in WT pups (Fig. 3A). Comparable root cell development was also observed in OV-TRβKO counterparts (Fig. 3B). The outer sulcus in OV-TRα/βKO mice, however, was largely composed of a single cuboidal epithelial layer bordering the endolymphatic space (Fig. 3C), suggesting a delay in the root cell differentiation. Consistent with the histological observations, OV-TRα/βKO showed fewer acetyl-α-tubulin positive cell processes within the spiral ligament compared with those in WT and OV-TRβKO counterparts (Fig. 3D-F). To further demonstrate root cell differentiation, we employed in situ hybridization (ISH) to investigate expression patterns of *Slc26a4* (also known as *Pendrin*) and *Epyc* (*Epiphycan*) mRNA. These genes are known to be abundantly and selectively expressed by root cells(Gu et al., 2020; Royaux et al., 2003; Xie et al., 2023).) Strong signals of *Slc26a4* (Fig. 3G-I) or *Epyc* (Supplementary Fig. S1A-C) mRNA could be detected deep inside the spiral ligament in both WT and OV-TRβKO cochleae, but in OV-TRα/βKO cochleae, expression domains of both molecules were adjacent to the luminal surface of the spiral ligament, consistent with a delay in the root cell development. These data suggest that root cell development is normally initiated in OV-TRβKO but disrupted in OV-TRα/βKO pups.

**Fig. 3.**
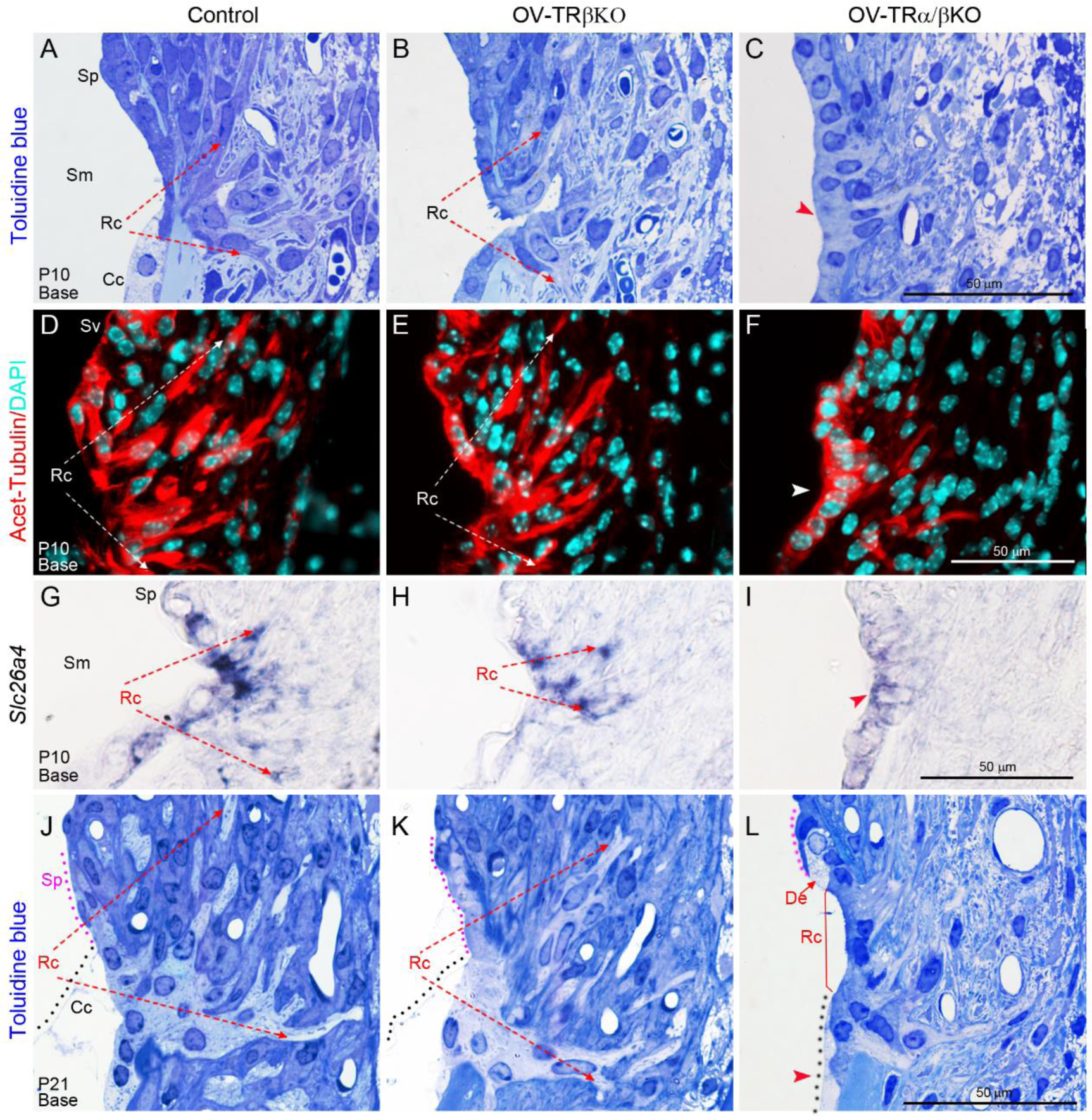
Otocyst-selective deletion of all known TRs retards root cell development. **(A-C)** Representative Toluidine Blue-stained sections showing delayed root cell (RC) development in OV-TRα/βKO pups at P10 (**C**), compared with that in control **(A)** or OV-TRβKO **(B),** Claudius cells (Cc), scala media (Sm), spiral prominence (Sp). **(D-F)** Immunostaining showing decreased acetyl-α-tubulin positive (red) root cell processes within OV-TRα/βKO outer sulcus **(F)**, compared with that in control **(D)** or OV-TRβKO counterparts **(E)** at P10. Nuclei were stained with DAPI (turquoise). n = 4 mice/ group. **(G-I)** ISH revealed a reduced expression domain of *Slc26a4* mRNA in OV-TRα/βKO (**I**), compared with that in control **(G)** or OV-TRβKO **(H)** at P10**. (J-L)** Representative sections showing disruption of root cell development in OV-TRα/βKO **(L),** compared with a mature outer sulcus in control **(J)** or OV-TRβKO **(K)** mice at P21. Cc is outlined by black dots, and Sp is outlined by magenta dots. A degenerated cell is indicated by “De” in (**L**). The upper basal turns were shown for all panels. Arrowheads indicate defects. Scale bars: 50 µm for all panels. n = 4 mice/group. See also Supplementary Fig. S1-2.

By P21, mature morphological features were comparable between OV-TRβKO and wild-type (WT) outer sulci, with elaborate root cell branches extending deep into the inferior region of the spiral ligament and upward behind the spiral prominence in the basal turn (Fig. 3J and K). Consistent with previous descriptions (Duvall III, 1969; Kimura, 1984), the surface of root cell bodies was covered by greatly enlarged Claudius cells and spiral prominence epithelium, no longer directly exposed to the endolymph at this stage. In contrast, the double knockout mice (OV-TRα/βKO) exhibited only initial signs of root cell differentiation at P21. Root cells with a few processes were mainly located along the scala media, separating the flattened Claudius cell and the spiral prominence epithelium, indicating a severe retardation in the differentiation of root cells and Claudius cells (Fig. 3L). Additionally, some cells with pale-staining cytoplasm and nucleus were occasionally detected in the OV-TRα/βKO outer sulcus at this age, suggesting degenerative changes (Fig. 3L). This observation was further supported by reduced acetyl-α-tubulin-positive root cell processes in the OV-TRα/βKO outer sulcus compared to WT or OV-TRβKO counterparts (Supplementary Fig. S1D-F).

By 1.5 months, the development of root cells and Claudius cells in OV-TRα/βKO mice had progressed to some extent but still lagged behind the developmental status observed in WT controls (Supplementary Fig. S2A-B, A’ and B’). Additionally, degenerative changes were noted in root cells and cochlear outer border cells (Boettcher’s cells) in some OV-TRα/βKO samples (Supplementary Fig. S2C and C’).

Taken together, these data demonstrate that root cell development is largely unaffected in OV-TRβKO mice but significantly delayed in OV-TRα/βKO mice. This suggests that while TRβ may not be individually indispensable, TRα and TRβ collectively play redundant roles in root cell development.

### Otocyst-selective deletion of all *TRs* causes early-onset root cell degeneration

Although the degeneration of cochlear hair cells and supporting cells has been reported previously in developmental hypothyroidism, and in mice with global deletion of TH transporters (Uziel et al., 1983; Sharlin et al., 2018; Mustapha et al., 2009), it remains unclear whether the survival of root cells depends on TH signaling. As described above, sporadic degenerative cells were occasionally detected in OV-TRα/βKO outer sulcus at P21 (Fig. 3L), or ∼1.5 months of age (Supplementary Fig. S2C and C’). By 2 months, WT control or OV-TRβKO outer sulcus showed no obvious degenerative changes, whereas OV-TRα/βKO exhibited reduced root cells with pale-staining processes and nuclei in both basal and apical turns (Fig. 4A-C, A’-C’, and Supplementary Fig. S3), suggesting progressive degenerations. Concomitant with the root cell degeneration, variable degenerations of the organ of Corti were observed in OV-TRα/βKO cochlea (Fig. 4C, Supplementary Fig. S3C), resembling those described previously in developmental hypothyroidism, or in mice with global deletion of TH transporters (Mustapha et al., 2009; Sharlin et al., 2018; Uziel et al., 1983).

**Fig. 4.**
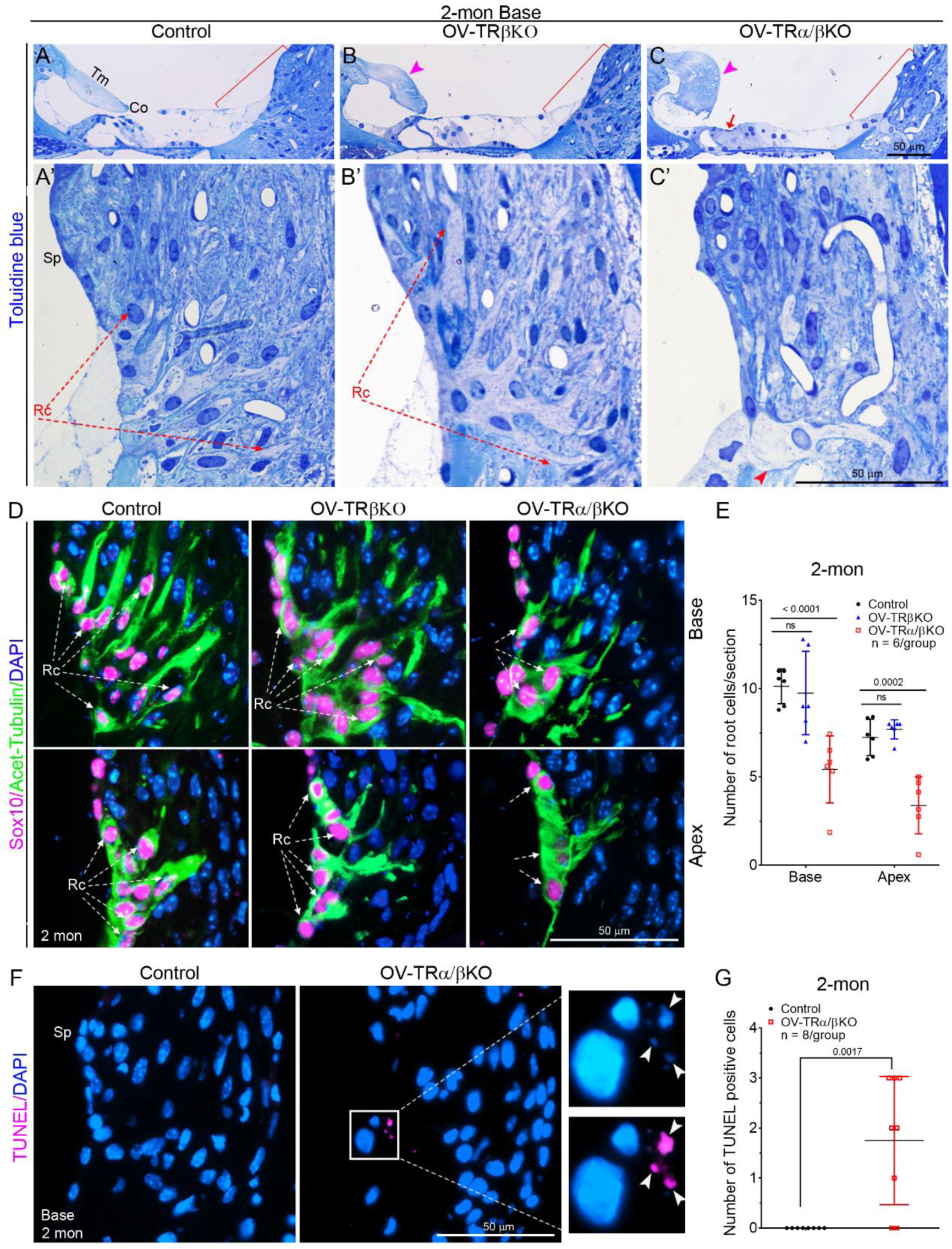
Otocyst-selective deletion of all TRs causes early-onset root cell degeneration. **(A-C)** Representative Toluidine Blue-stained sections of upper basal turn showing malformed tectorial membrane (Tm, Magenta arrowheads), a collapsed organ of Corti (Co, red arrow) in OV-TRα/βKO cochleae at 2 months **(C)**. **(A’)**, **(B’)** and **(C’)** are magnified views of outer sulcus regions indicated by red brackets in **(A)**, **(B)**, and **(C)**, respectively. OV-TRα/βKO showed reduced root cells with pale-staining processes (**C’**). Sp, Spiral prominence. n = 6 mice/group. **(D-E)** Representative sections (**D**) and a bar graph (**E**) show a decreased number of Sox10 (magenta) and acetyl-α-tubulin (green) double-positive root cells in OV-TRα/βKO outer sulcus, compared with that in control or OV-TRβKO at 2 mon. Nuclei were stained with DAPI (blue). N = 6 mice/group. (**F-G**) Representative sections (**F**) and a bar graph (**G**) show increased TUNEL-positive cells (magenta) in the OV-TRα/βKO outer sulcus at 2 mon, compared with that in the control group. The right column in (**F**) is a magnified view of the boxed area in the middle column, with separated color channels to better show the colocalization of TUNEL-positive signals with DAPI-stained (blue) nuclei (indicated by arrowheads). n = 8 mice/group. All values are represented as mean ±SD. A two-way ANOVA and an unpaired T-test were used to compare differences in (**E**) and (**G**), respectively. *P* Values were indicated. ns, not significant. Scale bars: 50 µm for all panels. See also Supplementary Fig. S3.

To assess root cell viability, we quantified the number of root cell nuclei at 2 months of age. Previous studies, including ours, have shown that Sox10 is localized in the nucleus of cochlear non-sensory cells, including outer border cells, root cells, and spiral prominence epithelium, but is excluded from fibrocytes in the lateral wall (Hao et al., 2014; Xie et al., 2023). Therefore, by performing double immunostaining for Sox10 and acetyl-α-tubulin (intensely expressed in the root cell cytosol), root cells can be easily identified as double-positive cells in the spiral ligament (Fig. 4D). Consistent with the histological observations, the number of root cell’s nucleus was significantly decreased in OV-TRα/βKO but not in OV-TRβKO, compared to that in WT (Fig. 4D-E). This supports the occurrence of root cell degeneration in the absence of both TRs.

Furthermore, we conducted a TUNEL analysis to detect apoptotic cells in the outer sulcus region. We observed sporadic yet limited TUNEL-positive cells in OV-TRα/βKO mice, but not in WT controls, at 2 months of age (Fig. 4F). Quantitative analysis revealed a statistically significant increase in TUNEL-positive cells in OV-TRα/βKO mice compared to the WT control group (Fig. 4G), suggesting apoptosis occurs at a low rate upon TRs loss. These data collectively indicate that TRα and TRβ function redundantly for root cell survival.

### Stria vascularis remains intact in OV-TRα/βKO mice

Given that OV-TRα/βKO mice exhibited a reduced EP compared to control mice (refer to Fig. 2E), we then examined the structural and functional integrity of SV, which plays an essential role in generating EP and maintaining K^+^ homeostasis in the endolymph(Hibino et al., 2010; Kim et al., 2013; Wangemann, 2006). At P21, no obvious histological abnormalities were observed in either OV-TRβKO or OV-TRα/βKO SV (Fig. 5A). The area of SV in both mouse lines was comparable to that in WT mice (Fig. 5B). Consistently, immunostaining analysis revealed normal expression patterns of EP-generating ion-transport molecules, including KCNJ10 (Kir4.1, a marker for intermediate cells)(Ando and Takeuchi, 1999; Marcus et al., 2002), Na^+^/K^+^-ATPase α(Kerr et al., 1982), NKCC1(Dixon et al., 1999) and KCNQ1 (Kv7.1)(Casimiro et al., 2001; Sakagami et al., 1991) (markers for marginal cells) (Fig. 5*C*). These data suggest that SV remains intact upon all *TRs* deleted in cochlear epithelium. Indeed, potential primary defects can be excluded from SV intermediate cells and basal cells in either OV-TRβKO or OV-TRα/βKO mice, as these cell types do not originate from the otic epithelium (Steel and Barkway, 1989) and thus should display no Cre recombination activity (refer to Fig. 2A).

**Fig. 5.**
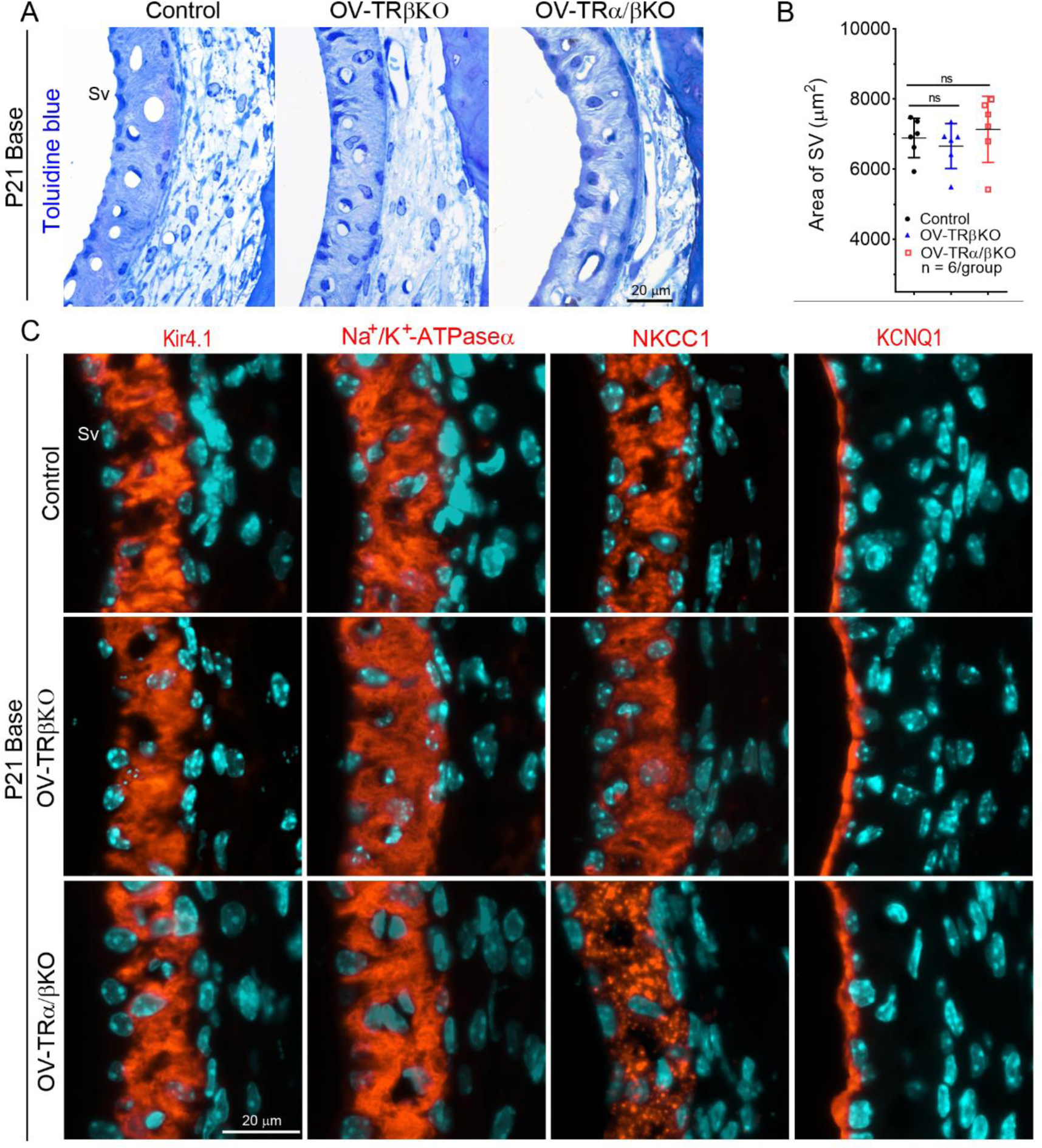
Stria vascularis (SV) remains intact in OV-TRα/βKO mice. (**A**) Representative Toluidine Blue-stained sections of cochlear upper basal turns showing comparable SV histology among Control, OV-TRβKO, and OV-TRα/βKO mice at P21. (**B**) Bar graph showing comparable SV areas among Control, OV-TRβKO, and OV-TRα/βKO mice at P21. n = 6 mice/group. Values are represented as mean ±SD. A one-way ANOVA followed by post-hoc pairwise tests (Bonferroni method) was used to compare differences between the two groups. ns: not significant. (**C**) Immunostaining revealing comparable expressions of Kir4.1, Na+/K+-ATPase α, NKCC1, and KCNQ1 (red) among Control, OV-TRβKO, and OV-TRα/βKO mice at P21. n = 3 mice/group for each molecular marker. Scale bars: 20 µm for all panels. Nuclei were stained with DAPI (turquoise). See also Supplementary Fig. S4.

As gap junctions are also essential for EP generation and hearing(Cohen-Salmon et al., 2002; Sun et al., 2005). We then analyzed the expression of Connexin26 (Cx26) and Cx30 proteins, which are the dominant gap-junction channel-forming proteins expressed in the cochlear epithelium(Ahmad et al., 2003). Comparable expression patterns of Cx26 and Cx30 were observed between control and OV-TRα/βKO cochleae at P21 (Supplementary Fig. S4). This suggests the gap junction system remains intact despite TRs deficiency.

### TH deficiency leads to the downregulation of a considerable number of root cell-enriched genes

To unravel the molecular mechanisms underlying outer sulcus development regulated by TH, we employed a microdissection technique to isolate outer sulcus regions from both euthyroid and hypothyroid mice at P10. Subsequently, we conducted a whole transcriptome analysis using RNA sequencing (RNA-seq). Each experimental group comprised five biological replicates. We identified 863 differentially expressed genes (DEGs) with a fold change of ≥ 2 and a Q-value of ≤ 0.05 (Supplementary Fig. S5, Supplementary Dataset S1). Genes with Fragments per kilobase per million mapped reads (FPKM) < 10 were excluded from the analysis.

Given that the microdissected region primarily consists of outer sulcus epithelial cells and fibrocytes, and our research focus is on root cell development, we conducted an analysis to identify DEGs preferentially expressed by root cells. This involved intersecting our dataset with a previously reported list of root cell-enriched genes (Jean et al., 2023). We identified a total of 178 root cell-enriched DEGs, consisting of 11 upregulated and 167 downregulated genes (Fig. 6A). Notably, several known marker genes for root cells, such as *Slc26a4* (Royaux et al., 2003), *Lgr5*, *Epyc* (Gu et al., 2020; Xie et al., 2023), and *P2rx2* (Lee et al., 2001), were among the downregulated DEGs identified, consistent with a delayed root cell differentiation in hypothyroid mice. Interestingly, 11 of the downregulated genes, including *Slc26a4*, *Tmprss3*, Gas2, *S100a1*, *Qsox1*, *Tmprss9*, *Prss36*, and *P2rx2*, have been associated with hearing loss in humans and/or mice when mutated (Groza et al., 2023) (https://hereditaryhearingloss.org; https://www.mousephenotype.org) (Fig. 6B).

**Fig. 6.**
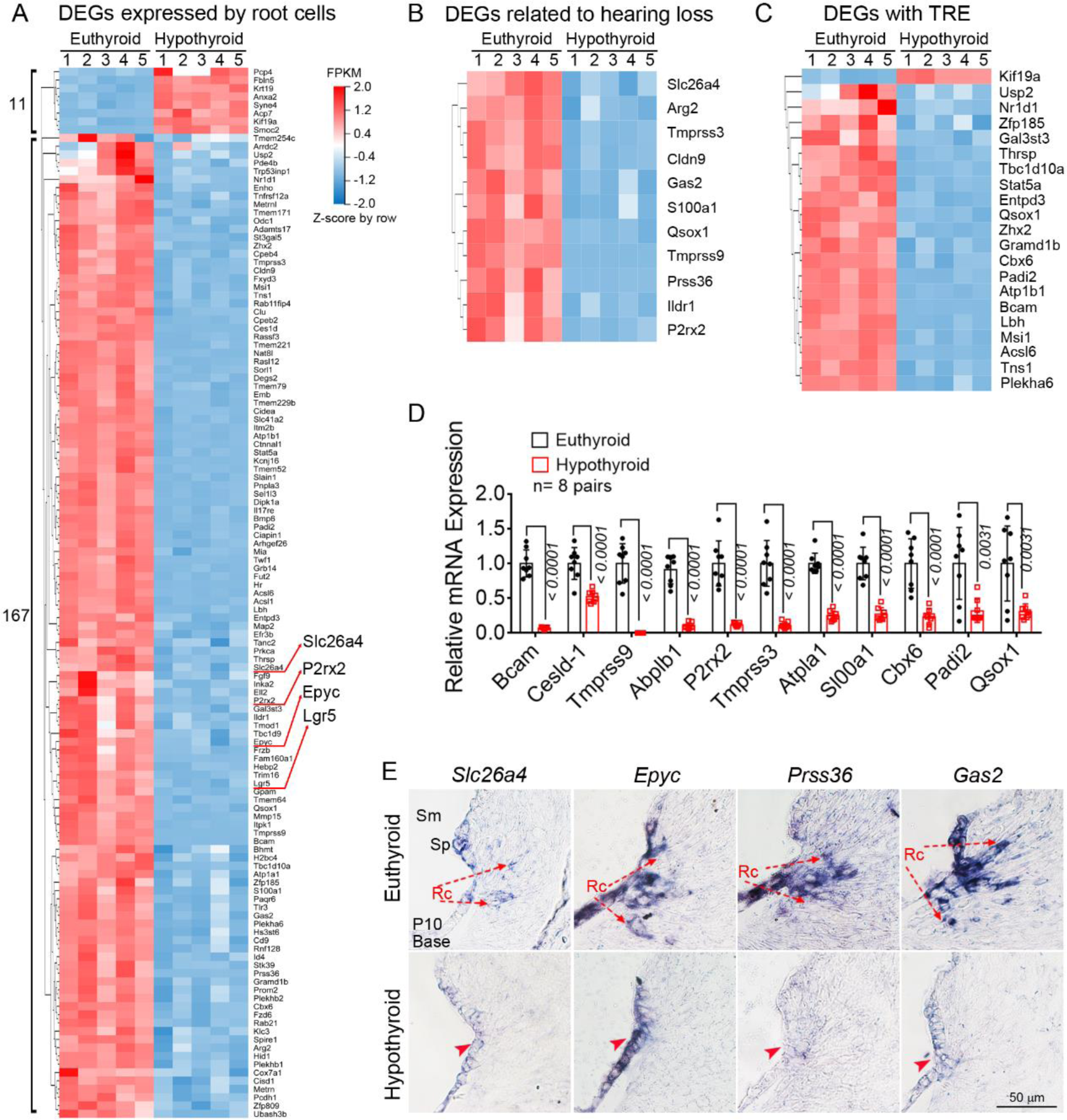
Deficiency of TH leads to the downregulation of a significant portion of root cell-enriched genes. **(A)** Heatmap displaying 11 upregulated and 167 downregulated DEGs preferentially expressed by root cells in hypothyroid compared with those in euthyroid mice at P10 revealed by RNA-seq (n = 5 pairs). **(B-C)** Heatmap displaying expression levels of 11 DEGs associated with hearing loss when mutated in humans or mice (**B**) and 21 DEGs carrying proximal TRE (**C**). (**D**) Validation of DEGs by qRT-PCR analysis. n = 8 pairs. Values are represented as mean ±SD. An unpaired T-test was used to compare differences. (**E**) Validation of DEGs by ISH analysis. Representative sections showing decreased mRNA expression domains of indicated genes at outer sulcus regions of hypothyroid mice (indicated by red arrowheads) compared with euthyroid counterparts at P10. n = 3 pairs. Rc: root cell, Sm: scala media, Sp: spiral prominence. Scale bar: 50 µm. See also Supplementary Fig. S5 and Dataset S1.

It is well established that the biological activities of TH are mainly mediated by TRs, which bind to TH response elements (TREs) to regulate target gene expression. The most frequently characterized TRE motif is a direct hexameric repeat of 5’(A/G)GGT(C/A)A3’ with 4 base spacer (called DR4 elements) (Chatonnet et al., 2013; Umesono et al., 1991). We, therefore, utilized the RcisTarget tool (https://bioconductor.org) to search for DEGs containing proximal TRE motifs to identify potential direct target genes of TH. We identified 21 potential direct target genes of TRs, with all but one showing downregulation in the hypothyroid outer sulcus (see Fig. 6C). Notably, among these genes, *Thrsp* has been previously established as a direct target gene of TRs in the liver (Zilz et al., 1990). Moreover, a significant number of these genes have been found to harbor endogenous TR binding sites in their proximal regions in the liver or nervous system, including *Thrsp, Usp2* (Grøntved et al., 2015), *Stat5a* (Chatonnet et al., 2013), *Nr1d1*, *Tbc1d10a*, *Qsox1*, *Zhx2*, *Atp1b1*, *Lbh*, *Tns1*, and *Plekha6* (Mendoza et al., 2021), supporting a potential direct regulation by TH. Further investigation is needed to determine if this also applies to the cochlea.

Fifteen DEGs were chosen for additional validation through qRT-PCR (Fig. 6D) or ISH (Fig. 6E). Selection criteria included significant fold changes or potential relevance to auditory function. Notably, the expression of *Prss36* in the cochlea has not been previously characterized. Together, these data underscore the significance of TH signaling in regulating the gene expression profile associated with root cell differentiation.

## Discussion

This study highlights the pivotal role of TH and TRs in postnatal root cell development and survival. The absence of both TRα and TRβ, but not TRβ alone, in cochlear epithelium led to significant retardation in root cell differentiation and early-onset root cell degeneration. These findings suggest that TRα and TRβ function redundantly in promoting late-stage root cell differentiation and maintaining cellular homeostasis.

The critical period when TH is essential for cochlear development and normal hearing covers from late embryonic stages to around the second postnatal week in rodents(Deol, 1973b; Uziel et al., 1985; Uziel, 1986). During this developmental window, the cochlear sensory epithelium undergoes dramatic morphological changes including regression of GER leading to the formation of the inner sulcus, and opening of the tunnel of Corti (Deol, 1973b; Roth and Bruns, 1992). Concomitantly, the outer sulcus epithelium transforms from a simple layer of cuboidal-shaped cells lining the scala media into root cells with large, branched processes residing deep in the spiral ligament(Lim and Anniko, 1985; Roth and Bruns, 1992; Xie et al., 2023). This study uncovers that TH or TRs are necessary for the outer sulcus remodeling, in addition to previously reported GER regression and the organ of Corti maturation(Rusch et al., 2001). Although both TRα and TRβ are expressed by the developing outer sulcus(Bradley et al., 1994; Gu et al., 2020; Jean et al., 2023; Ng et al., 2015), severe retardation in root cell differentiation was observed only in the absence of both receptors indicating that TRα and TRβ compensate functionally for each other in controlling root cell development. Our findings extend those data obtained from the mouse model with global deletion of TRs, which suggest synergistic effects of TRα and TRβ in controlling the remodeling of the organ of Corti and the correct formation of the tectorial membrane (Rusch et al., 2001).

Another intriguing finding of this study is the early-onset progressive degeneration of root cells observed only in OV-TRα/βKO but not OV-TRβKO cochlea, suggesting TRα and TRβ play a combined role in root cell survival. Interestingly, degeneration of root cells and the organ of Corti initiated concomitantly in juvenile OV-TRα/βKO cochlea, indicating potential coordinated mechanisms underlying these degenerations. As root cells, with their extensive processes, are closely associated with capillaries and fibrocytes within the spiral ligament, it has been proposed that they may provide a cellular route for transporting nutrients, blood-borne hormones (such as T3), metabolites, and ions between the vasculature in the spiral ligament into the inferior cochlear sensory epithelium(Duvall III, 1969; Jagger and Forge, 2013; Shodo et al., 2017), which lacks direct blood flow. This hypothesis is supported by an epithelial gap junction network formed by the syncytium of supporting cells in the organ of Corti and extending through the Claudius’ cells and root cells(Jagger and Forge, 2015, 2006). Hence, it’s reasonable to assume that the proper timely development of root cells may contribute to maintaining the health of the cochlear sensory epithelium. The early-onset degeneration of root cells might compromise the supply of nutrients and hormones or disrupt ion and metabolite homeostasis in the organ of Corti, thereby promoting its degeneration. Alternatively, the health of root cells might be influenced by the degeneration of the organ of Corti. Severe early-onset degenerations of cochlear hair cells have been reported previously in congenital hypothyroidism or mice with global deletion of thyroid hormone transporters (Mustapha et al., 2009; Sharlin et al., 2018), although the status of root cells in these animal models has not been described. It would be valuable to investigate any alterations in root cell structure or function in these mice.

Several lines of evidence including our recently published report suggest that root cells may play a significant role in the regulation of endolymph homeostasis and EP (Spicer and Schulte, 1996; Jagger et al., 2010; Cazals et al., 2015; Xie et al., 2023). For instance, the absence of the KCNK5 channel, primarily located in the outer sulcus, leads to severe hearing loss accompanied by a more than 60% decrease in EP magnitudes (Cazals et al., 2015). Furthermore, our recent study revealed that the disruption of outer sulcus development due to *Zbtb20* deletion results in a reduction of over 50% in EP values (Xie et al., 2023). In this study, however, OV-TRα/βKO mice showed only a reduction of approximately 13% and 22% in mean EP values at 1.5 months and 4 months, respectively. This discrepancy could be attributed to varying degrees of disruption in root cell development and function among different animal models. Unlike *Zbtb20* knockout mice, where there is a permanent disruption in outer sulcus differentiation, the outer sulcus development in OV-TRα/βKO mice had progressed to some extent by approximately 1.5 months. Subsequently, root cells experienced gradual degeneration, but approximately 50% of them persisted in OV-TRα/βKO mice. The mild EP reduction in OV-TRα/βKO mice may indicate a robust compensatory ability of root cells in regulating endolymph homeostasis and EP. Further investigation through targeted ablation of root cells could provide insight into the extent of their contribution to EP magnitude and clarify their role in regulating cochlear function.

It is noteworthy that the reduction in EP observed in OV-TRα/βKO mice is less pronounced than that documented in global *Thra* and *Thrb* double knockout mice (Rusch et al., 2001). This discrepancy indicates that TRs expression outside of the cochlear epithelium, particularly in spiral ligament fibrocytes, may also contribute to EP maintenance. This hypothesis is supported by the postnatal expression of both *Thra* and *Thrb* mRNA by fibrocytes (Jean et al., 2023), which are known to play a role in EP maintenance (Adachi et al., 2013; Furness, 2019). Our initial observations indicate a disruption in fibrocyte development in hypothyroid mice. However, in OV-TRα/βKO mice, primary fibrocyte defects can be ruled out as Cre recombination activity was confined in the otocyst-derived cochlear epithelium, as shown in Fig. 2A.

Although SV defects have been observed in cases of secondary hypothyroidism caused by pituitary transcription factor 1 (Pit1) mutation (Mustapha et al., 2009), there have been no reported SV alterations in cases of deficiency in TH transporters (Sharlin et al., 2018), or in global *TRs* double knockout mice(Rusch et al., 2001), both of which exhibit a significant reduction in EP. Consistently, our findings in this study show normal SV upon deletion of both *Thra* and *Thrb* in cochlear epithelium. Furthermore, previous studies utilizing a sensitive *LacZ* gene reporter system have shown that *Thrb* mRNA is not expressed in the developing SV (Ng et al., 2015). Therefore, the SV is unlike a primary defect site responsible for a reduced EP, which occurred only in mice lacking all TRs(Rusch et al., 2001).

In summary, this study highlights the redundant role of TH receptors (TRα and TRβ) in promoting the late-stage differentiation and survival of root cells, likely through the activation of a significant portion of genes enriched in root cells. These findings suggest that the expression of TRs in the cochlear epithelium is essential for maintaining an optimal EP magnitude and that TRs expressed in areas outside of the cochlear epithelium, such as spiral ligament fibrocytes, may also play a significant role in EP maintenance. These findings advance our understanding of how TH regulates cochlear remodeling and maturation and provide new insights into the mechanisms governing the development and function of cochlear outer sulcus regions.

## Materials and Methods

### Animal models

All animal maintenance and experimental procedures were conducted in compliance with the guidelines issued by the Animal Ethics Committee of Naval Medical University. Protocols (XIEC-NSFC-2022-033) were approved by the Ethics Committee of Xinhua Hospital affiliated with Shanghai Jiaotong University School of Medicine. *Thra* ^flox^(Liu et al., 2022), *Thrb*^flox^ (Yan et al., 2022), or ROSA26-*LacZ* reporter (Soriano, 1999)mice have been described previously. The OV-TRβKO mice were generated by cross-breeding of *Thrb* ^flox^ mice with *Foxg1*-*Cre* mice (Jax, Stock No. 004337)(Hebert and McConnell, 2000). The OV-TRα/βKO mice were then generated by cross-breeding of heterozygous OV-TRβKO mice with *Thra* ^flox^ mice. All mice were maintained on a C57BL/6J background. Floxed/*Cre*-negative littermate mice served as wild-type controls. PCR primers for genotyping are listed in Supplementary Table S1. To induce hypothyroidism, timed-pregnant C57BL/6J mice were treated with antithyroid agents (0.05% methimazole, 1.0% sodium perchlorate monohydrate, and 5.0% sucrose in drinking water) (Sawant et al., 2015) or vehicle from embryonic day 17. Treatment continued during lactation and ceased on the designated day. Data from both genders were pooled together because no gender differences in the phenotypes were observed.

### Histological analysis

Mice were transcardially perfused with 4% paraformaldehyde (pH 7.4) to fix the tissues. Cochleae were then dissected and perilymphatically perfused with 2.5% glutaraldehyde, followed by overnight fixation. Post-fixation was performed with 1% osmium tetroxide. Subsequently, the cochleae were decalcified, dehydrated, and embedded in Spurr’s low-viscosity resin. Mid-modiolar sections (1 µm) were stained with 1.25% (w/v) toluidine blue.

For quantitative analysis, images of the upper basal turns from three non-consecutive sections were captured at 200x magnification. The stria vascularis (SV) areas were measured using the Image-Pro Plus 6.0 program. Each analysis included a single cochlea from each animal. The number of animals analyzed at a given age was indicated in figure legends.

### Immunohistochemistry and X-Gal staining

Immunostaining was conducted following previously established protocols(Xie et al., 2023). X-gal staining was performed according to published methods (Xie et al., 2008). Details regarding primary antibodies are provided in Supplementary Table S2. To quantify root cells, non-consecutive mid-modiolar sections per cochlea underwent double immunostaining for Sox10 and acetyl-α-tubulin. The nuclei of double-positive cells within the outer sulcus region were manually counted from at least 5 sections for each animal, with the genotype blinded during counting. A single ear was analyzed for each animal.

### Auditory brainstem response (ABR) test

Auditory brainstem responses (ABRs) were recorded using a TDT RZ6/BioSigRZ system (Tucker-Davis Technologies Inc., USA), following established procedures (Xie et al., 2023). Briefly, anesthetized mice were placed on a heated mat within a sound-proof chamber. Click or pure tone stimuli ranging from 4 to 32 kHz were generated. The sound intensity was gradually decreased from 90 to 10 dB sound pressure level (SPL) in 5 dB steps, and auditory thresholds were determined as the lowest sound intensity that elicited reproducible and recognizable waves.

### Endocochlear potential (EP) measurement

EP recording via the cochlear lateral wall was conducted following previously described methods (Li et al., 2020; Xie et al., 2023). Both ears’ basal turns were analyzed for each animal. In brief, mice were anesthetized, and a tracheotomy was performed. The tympanic bulla was opened to expose the cochlea. A small hole was drilled in the cochlear lateral wall of the basal turn, through which a micropipette electrode filled with 150 mM KCl was carefully advanced into the scala media using a micromanipulator. The voltage changes during the insertion were continuously recorded using an amplifier (MultiClamp 700B, Molecular Devices) combined with a digitizer (Digidata 1440A, Molecular Devices). The EP magnitude of each ear was recorded and presented individually on the graph.

### In situ hybridization (ISH)

ISH was carried out according to previously published methods(Xie et al., 2023). In brief, mice were transcardially perfused with 4% paraformaldehyde (pH 9.0) and cochleae were dissected and perfused perilymphatically with the same fixative. The fixed and decalcified cochleae were cryostat sectioned (14 μm thickness). Sections were then subjected to ISH as described previously(Xie et al., 2010). Information on antisense cRNA probes is listed in Supplementary Table S3. Cochleae from n ≥ 4 mice per genotype at a given age were analyzed.

### TUNEL (Terminal deoxynucleotidyl transferase dUTP Nick-End labeling) assay

Semi-serial cochlear sections from mice at indicated ages were digested with 10 μg/mL proteinase K for 30 minutes at 37°C, followed by incubation with a TUNEL reaction mixture at 37°C for 2 hours. Images of the outer sulcus regions spanning from the basal to apical turns were captured at 400x magnification. The TUNEL positive nuclei were manually counted on 8 non-consecutive sections per ear. Analysis was conducted on one ear per animal.

### Real-time RT-PCR (qRT-PCR)

Total RNA extraction, cDNA synthesis, and qRT-PCR were performed as described previously(Xie et al., 2023). Data were normalized to 36B4 (Rplp0, internal control) and presented as fold changes of gene expression in the mutant mice compared to the WT Control. Each sample was analyzed in duplicate. The primers used are shown in Supplementary Table S1.

### Outer sulcus region isolation and RNA-seq analysis

To isolate the outer sulcus regions from either euthyroid or hypothyroid C57BL/6J mice at P10, the spiral ligament and attached basilar membrane were isolated in cold Hanks’ balanced salt solution buffer. Subsequently, the outer sulcus region was microdissected by carefully removing the adjacent basilar membrane and the reddish stria vascularis, along with the underlying spiral ligament, using two 1 ml syringes equipped with 26G needles. This dissection method ensured that the isolated subregion primarily contained the outer sulcus with its underlying fibrocytes, as well as the directly connected spiral prominence epithelium and Claudius cells.

The isolated outer sulcus regions from four ears were pooled together, and total RNA was extracted using the Qiagen RNeasy Micro kit. RNA-seq experiments were conducted with five biological replicates for each of the two conditions, following a previously established protocol(Xie et al., 2023). Clean reads obtained were aligned to the Mus musculus reference genome (GCF_000001635.26_GRCm38.p6) using HISAT2 (v2.0.4). Significant differentially expressed genes (DEGs) between euthyroid and hypothyroid outer sulcus samples were identified using DESeq2 (v1.4.5), applying criteria of |log2FC|≥1 and Q-value≤0.05, and filtering out genes with FPKM<10 in both conditions. DEGs that were preferentially expressed by root cells were identified by intersecting the DEG dataset with the top 800 root cell-enriched genes at P12, as identified in a previous study using single-cell RNA-seq (Jean et al., 2023).

To identify DEGs containing proximal TH response elements (TREs), the region spanning 10 Kb up and downstream of the transcription start site for each DEG (with reference to the Mus musculus mm10 genome) was scanned with a classical TRE motif (<u>THA_MOUSE.H11MO.0.C</u>, <u>THB_MOUSE.H11MO.0.D)</u>, which is accessible at https://hocomoco11.autosome.org. The RcisTarget tool 1.14.0, available at https://bioconductor.org, was employed for this analysis. The TRE employed is the most extensively characterized TRs binding consensus sequence. It comprises a direct hexameric repeat of 5’(A/G)GGT(C/A)A3’ with a 4-base spacer, commonly referred to as DR4 elements (Chatonnet et al., 2013; Umesono et al., 1991).

### Statistical analysis

All statistical analyses were conducted using GraphPad Prism 7 software. Data normality tests were initially performed. Pairwise comparisons were performed using either an unpaired Student’s t-test or a non-parametric Mann-Whitney test. For multiple group comparisons, a Kruskal-Wallis test (a non-parametric alternative to ANOVA) with Dunn’s multiple comparisons test or a two-way ANOVA followed by post-hoc pairwise tests (Bonferroni method) was utilized when applicable. Values are presented as mean ± SD or as medians with interquartile ranges (25-75%) for non-parametric tests exclusively. Tests and groups are noted in figure legends. In all instances, a two-tailed p-value < 0.05 was considered statistically significant. Non-significant differences between samples are denoted as “ns,” while p-values for significant differences are displayed in the graphs.

## Author Contributions

ZX and WZ conceived the idea, and wrote the manuscript; ZX, WZ, DH, and JT analyzed data and revised the manuscript; HY provided animal models; XM, FJ, JT, LS, YC, and SH performed most of the experiments. CW analyzed RNA-seq data. All authors were involved in the final approval of the submission of the manuscript.

## Acknowledgments and fundings

This work was supported by grants from the National Key R&D Program (2019YFA0802500, 2018YFA0800602), the National Natural Science Foundation of China (32271162), and the Collaborative Innovation Program of Shanghai Municipal Health Commission (2020CXJQ01).

## Data availability

RNA-seq data are accessible in Gene Expression Omnibus (GEO) under the accession number <u>GSE268897</u> with the hyperlinks https://www.ncbi.nlm.nih.gov/geo/query/acc.cgi?acc=GSE268897 (Xie, et al, 2024). Other data supporting the findings of this study are available within the manuscript and its Supplementary information files.

## Supplementary information

**Table S1.**
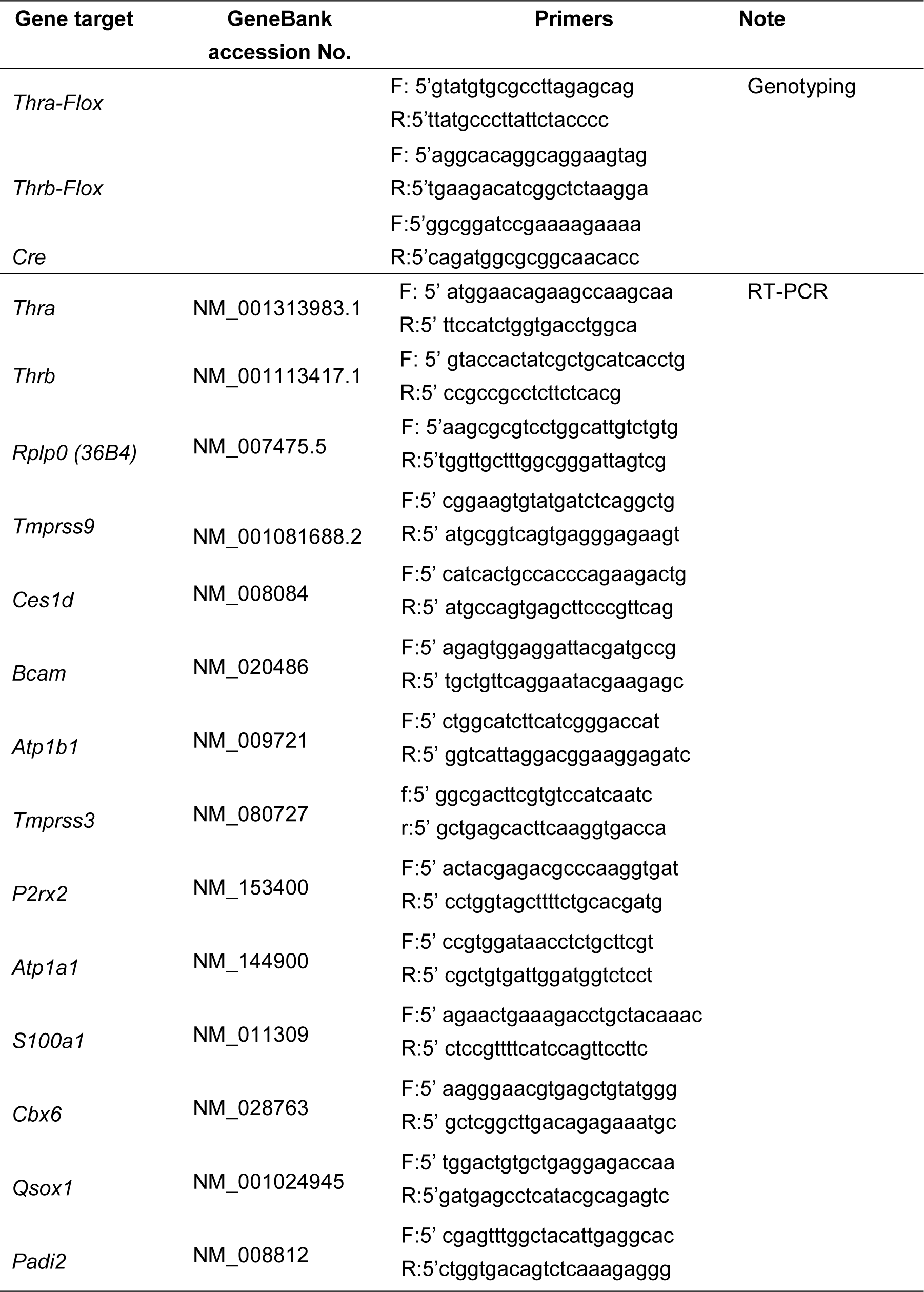
Sequence information for PCR primers.

**Table S2.** The information for primary antibodies.

| **Antibody** | **Resource** | **Dilution** |
| --- | --- | --- |
| anti-alpha 1 Sodium<br>Potassium ATPase | Abcam, ab7671 | 1:2000 |
| anti-KCNQ1 | Santa Cruz, SC-10646 | 1:2000 |
| anti-Kir4.1 | Alomone labs, APC-035 | 1:2000 |
| anti-NKCC1 | Invitrogen, PA5-98154 | 1:2000 |
| anti-Acetylated Tubulin | Sigma, T7451 | 1:300 |
| anti-Sox10 | Abclonal, A15100 | 1:2000 |
| anti-CX26 | Abcam, ab65969 | 1:2000 |
| anti-CX30 | Invitrogen, 33-2500 | 1:2000 |

**Table S3.**
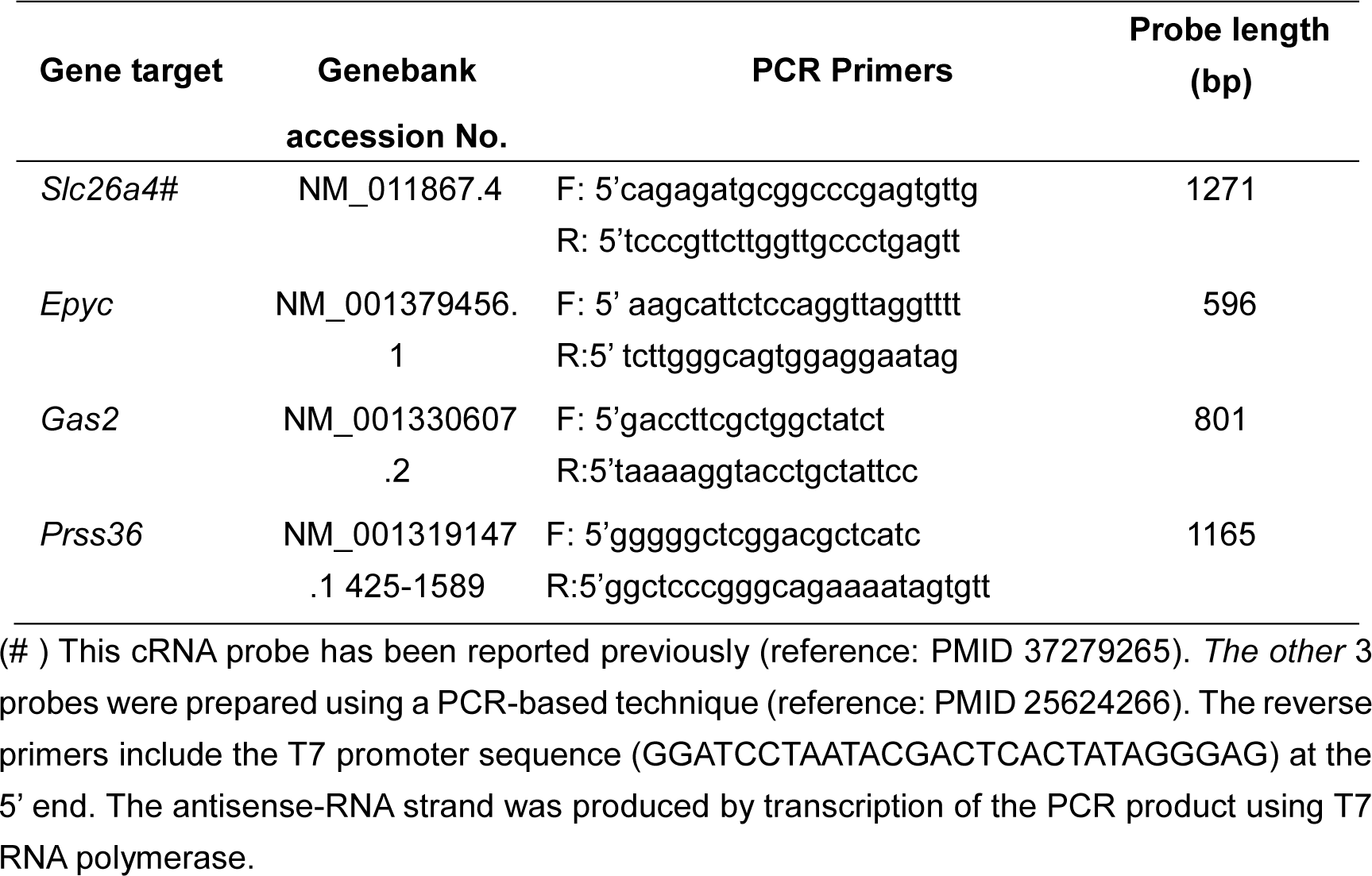
Sequence information for cRNA probes.

**Fig. S1.**
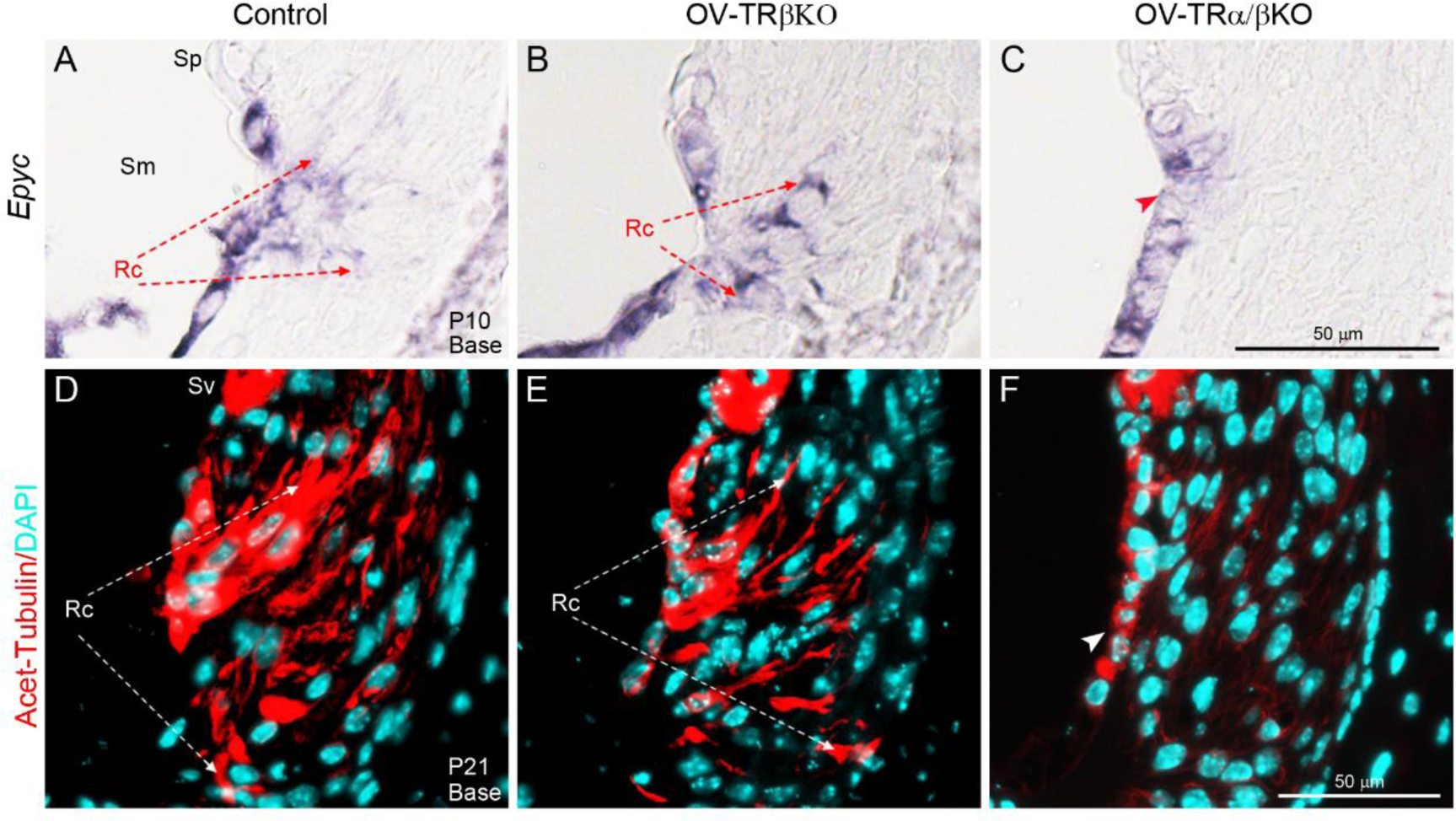
Cochlea-selective deletion of all known TRs retards root cell development. **(A-C)** ISH showing a decreased expression domain of *Epyc* mRNA (a root cell marker) in OV-TRα/βKO outer sulcus **(C)** at P10, compared with that in control **(A)** or OV-TRβKO **(B)** counterparts. n = 4 mice/group. **(D-F)** Representative immunostained sections showing reduced acetyl-α-tubulin positive (red) root cell processes within the inferior region of the spiral ligament from OV-TRα/βKO basal turns **(F)** compared with those from control **(D)** or OV-TRβKO counterparts **(E)** at P21. Scale bars: 50 µm for all panels. Nuclei were stained with DAPI (turquoise). n = 4 mice/group. Rc, root cell; Sm, scala media; Sv, stria vascularis; Sp, spiral prominence. Related to Fig.3.

**Fig. S2.**
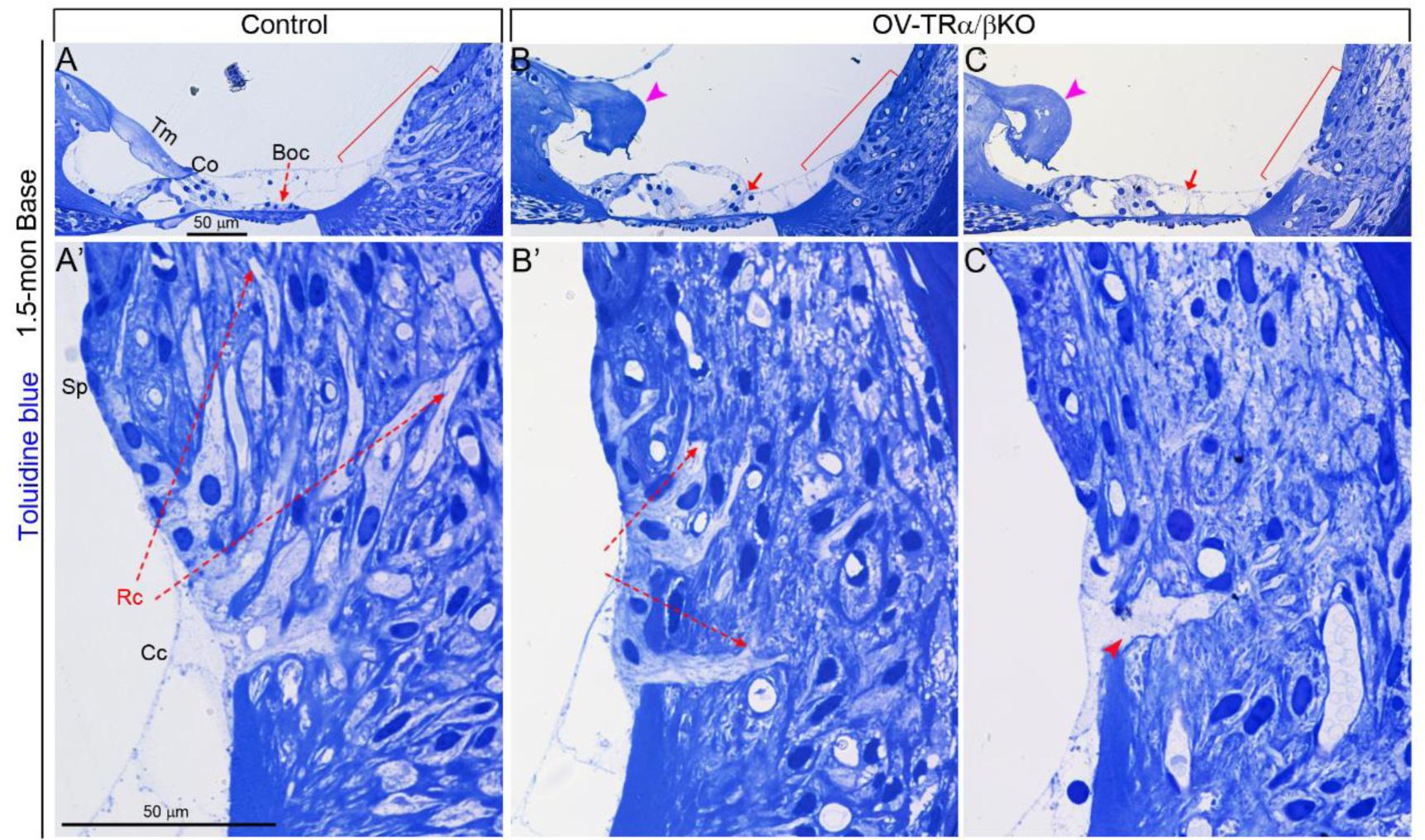
Development retardation compound with degeneration in root cell from OV-TRα/βKO mice at 1.5 months of age. **(A-C)** Representative Toluidine Blue-stained sections of upper basal turns from control **(A)**, and OV-TRα/βKO **(B and C)** cochleae at 1.5 months of age. Magenta arrowheads in **(B and C)** indicate tectorial membrane (Tm) malformations in OV-TRα/βKO cochleae. Red arrows indicate either delayed differentiation **(B)** or degeneration **(C)** of Boettcher cells (Boc). **(A’)**, **(B’)** and **(C’)** are magnified views of outer sulcus regions indicated by red brackets in **(A)**, **(B)**, and **(C)**, respectively. **(A’)** Control mice showed numerous branched root cells (Rc) processes extending deep into the inferior region of the spiral ligament and upward to behind the spiral prominence (Sp), **(B’)** whereas OV-TRα/βKO displayed reduced root cell processes within the superficial region of the spiral ligament. **(C’)** Some samples from OV-TRα/βKO mice showed degenerative changes, displaying reduced root cells with pale-staining processes (indicated by red arrowhead). Cc, Claudius cells; Spiral prominence. Scale bars: 50 µm for all panels. n = 4 mice/group. Related to Fig. 3.

**Fig. S3.**
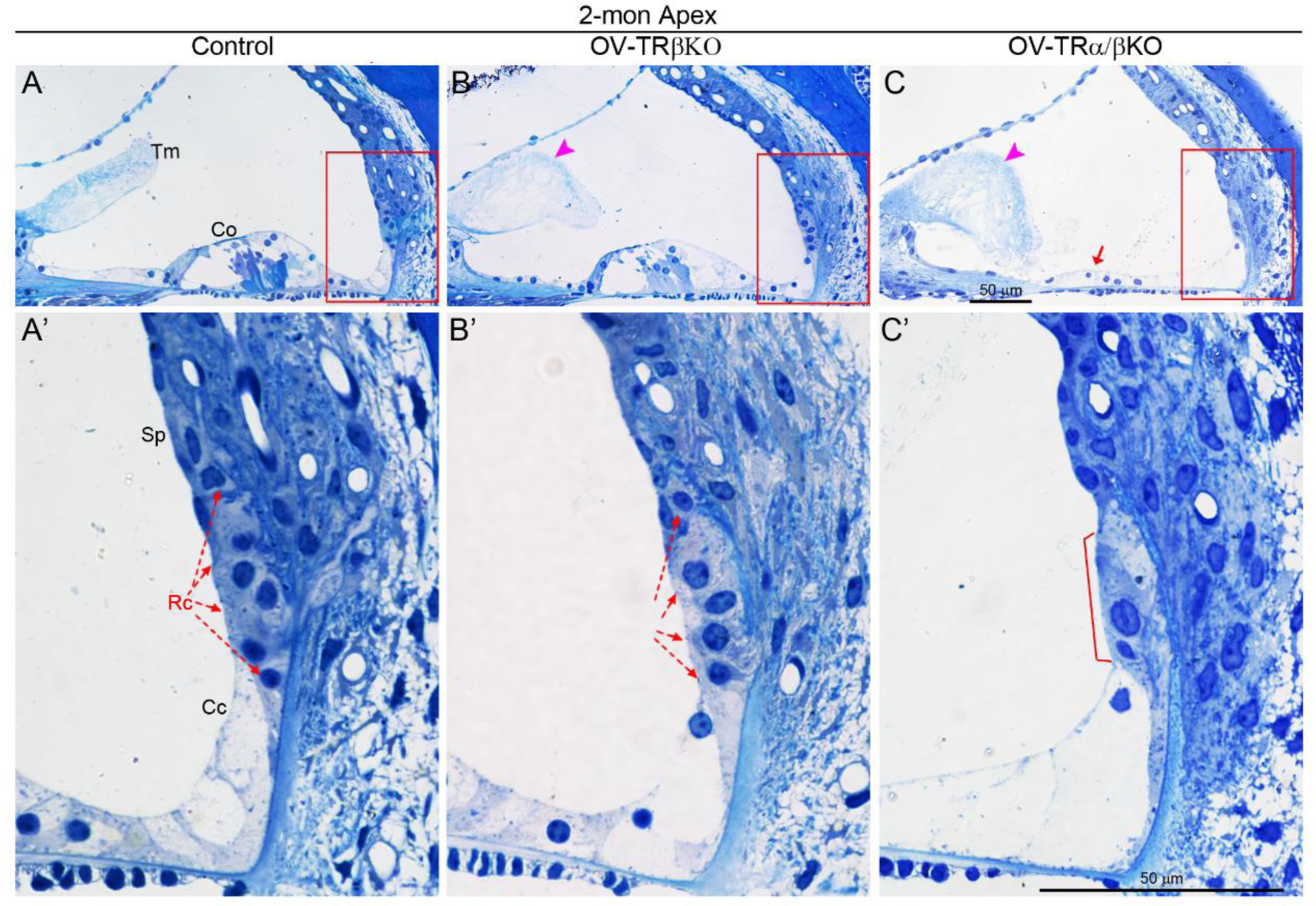
Absence of all known TRs in cochlear epithelium causes early-onset root cell degeneration. **(A-C)** Representative Toluidine Blue-stained sections of apical turns show an upright organ of Corti (Co) in control **(A)** or OV-TRβKO **(B)**, whereas only sparse cells (indicated by the red arrow) in OV-TRα/βKO epithelium **(C)** at 2 months of age. Tectorial membrane (Tm, indicated by magenta arrowheads) malformations were observed in both OV-TRβKO and OV-TRα/βKO. **(A’)**, **(B’)** and **(C’)** are magnified views of boxed region in **(A)**, **(B)**, and **(C)**, respectively, showing reduced root cells (Rc) in OV-TRα/βKO outer sulcus (indicated by a red bracket) **(C’)**, compared with those in control **(A’)** or OV-TRβKO **(B’)**. Sp, Spiral prominence. n = 6 mice/group. Scale bars: 50 µm for all panels. Related to Fig. 4.

**Fig. S4.**
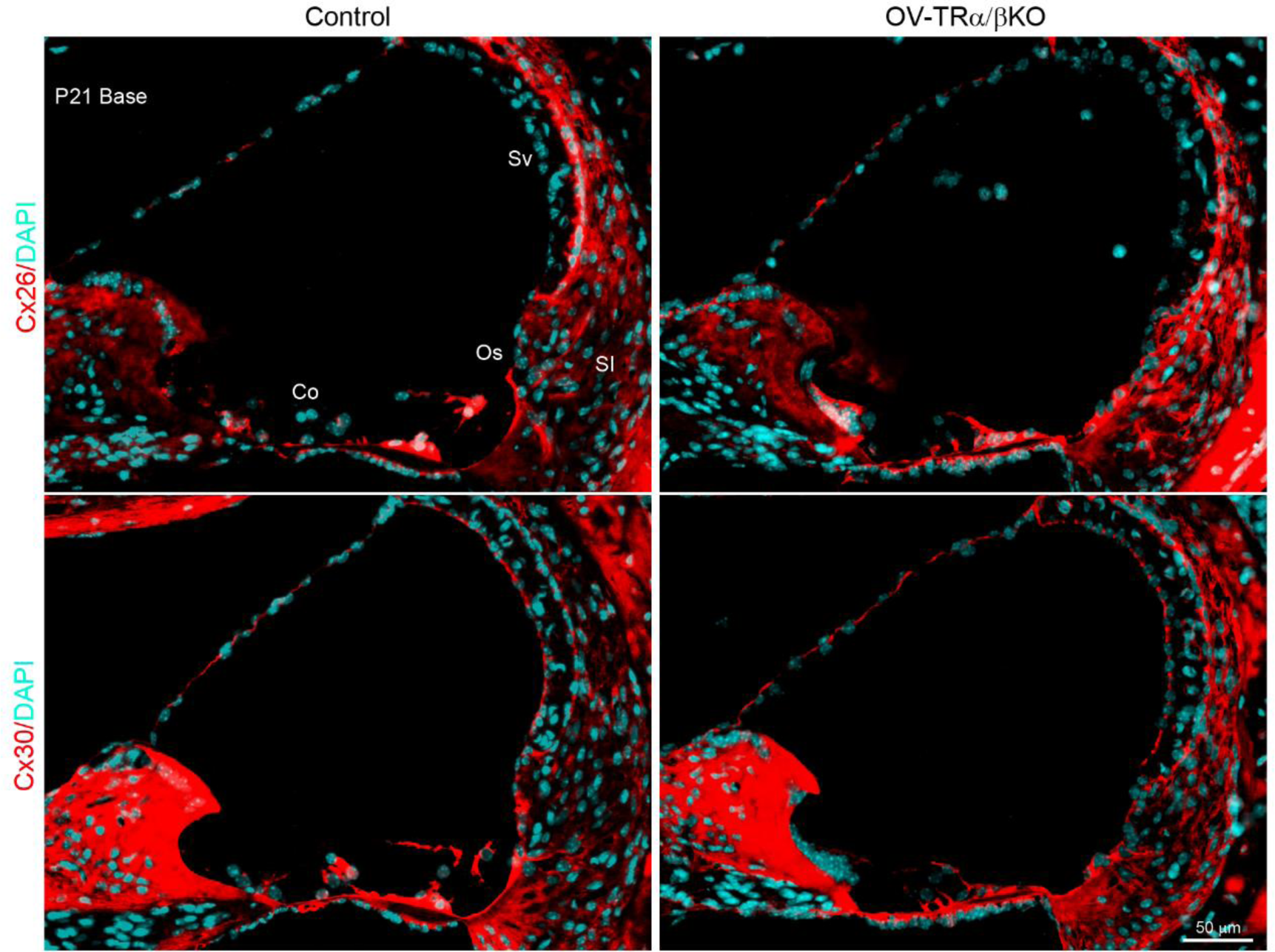
Ablation of TRs does not affect the expression patterns of connexins in cochleae. Representative immuno-stained sections of upper basal turn showing comparable expression patterns of CX26 and CX30 (red) between Control and OV-TRα/βKO mice at P21. Nuclei were stained with DAPI (turquoise). Co, organ of Corti; Os, outer sulcus; Sl, spiral ligament; Sv, stria vascularis. Scale bars: 50 µm. n = 3 mice/group. Related to Fig. 5.

**Fig. S5.**
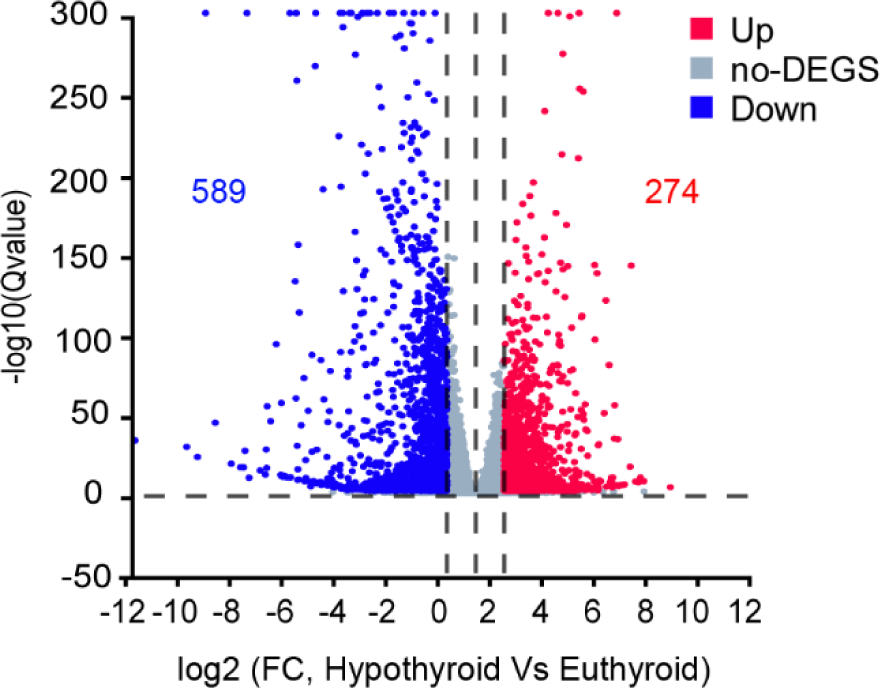
Volcano plot showing the significantly differentially expressed genes (DEGs) detected in microdissected outer sulcus regions from hypothyroid cochlea at P10 compared to euthyroid counterparts by RNA-seq analysis (n = 5 pairs, fold change ≥ 2, Q value ≤ 0.05). Related to Fig. 6 and Supplementary Dataset 1.

**Dataset S1 (separate file):** List of total DEGs and DEGs preferentially expressed by root cells in microdissected outer sulcus regions from hypothyroid cochlea at P10 compared to euthyroid counterparts by RNA-Seq analysis. DEGs were identified based on a fold change ≥ 2, Q value ≤ 0.05 (Genes of FPKM (Fragments per kilobase per million mapped reads) < 10 were filtered out). Related to Fig. 6 and Supplementary Fig. S5.

## Reference

Xie Z, Zhang W. Gene expression profiles of microdissected cochlear outer sulcus regions from eu- and hypothyroid mice at P10. GEO. https://www.ncbi.nlm.nih.gov/geo/query/acc.cgi?acc=GSE268897. Deposited 3 Jun 2024.

Adachi N, Yoshida T, Nin F, Ogata G, Yamaguchi S, Suzuki T, Komune S, Hisa Y, Hibino H, Kurachi Y. 2013. The mechanism underlying maintenance of the endocochlear potential by the K+ transport system in fibrocytes of the inner ear. J Physiol 591:4459–4472. doi:10.1113/jphysiol.2013.258046

Ahmad S, Chen S, Sun J, Lin X. 2003. Connexins 26 and 30 are co-assembled to form gap junctions in the cochlea of mice. Biochemical and Biophysical Research Communications 307:362–368. doi:10.1016/S0006-291X(03)01166-5

Ando M, Takeuchi S. 1999. Immunological identification of an inward rectifier K+ channel (Kir4.1) in the intermediate cell (melanocyte) of the cochlear stria vascularis of gerbils and rats. Cell Tissue Res 298:179–183. doi:10.1007/s004419900066

Bradley DJ, Towle HC, Young WS. 1994. Alpha and beta thyroid hormone receptor (TR) gene expression during auditory neurogenesis: evidence for TR isoform-specific transcriptional regulation in vivo. Proc Natl Acad Sci U S A 91:439–443. doi:10.1073/pnas.91.2.439

Casimiro MC, Knollmann BC, Ebert SN, Vary JC, Greene AE, Franz MR, Grinberg A, Huang SP, Pfeifer K. 2001. Targeted disruption of the Kcnq1 gene produces a mouse model of Jervell and Lange-Nielsen Syndrome. Proc Natl Acad Sci U S A 98:2526–2531. doi:10.1073/pnas.041398998

Cazals Y, Bevengut M, Zanella S, Brocard F, Barhanin J, Gestreau C. 2015. KCNK5 channels mostly expressed in cochlear outer sulcus cells are indispensable for hearing. Nat Commun 6:8780. doi:10.1038/ncomms9780

Chatonnet F, Guyot R, Benoît G, Flamant F. 2013. Genome-wide analysis of thyroid hormone receptors shared and specific functions in neural cells. Proc Natl Acad Sci U S A 110:E766–775. doi:10.1073/pnas.1210626110

Christ S, Biebel UW, Hoidis S, Friedrichsen S, Bauer K, Smolders JWT. 2004. Hearing loss in athyroid pax8 knockout mice and effects of thyroxine substitution. Audiol Neurootol 9:88–106. doi:10.1159/000076000

Cohen-Salmon M, Ott T, Michel V, Hardelin J-P, Perfettini I, Eybalin M, Wu T, Marcus DC, Wangemann P, Willecke K, Petit C. 2002. Targeted Ablation of Connexin26 in the Inner Ear Epithelial Gap Junction Network Causes Hearing Impairment and Cell Death. Current Biology 12:1106–1111. doi:10.1016/S0960-9822(02)00904-1

Deol MS. 1973a. Congenital deafness and hypothyroidism. Lancet 2:105–6. doi:10.1016/s0140-6736(73)93310-2

Deol MS. 1973b. An Experimental Approach to the Understanding and Treatment of Hereditary Syndromes with Congenital Deafness and Hypothyroidism. Journal of Medical Genetics 10:235. doi:10.1136/jmg.10.3.235

Dixon MJ, Gazzard J, Chaudhry SS, Sampson N, Schulte BA, Steel KP. 1999. Mutation of the Na-K-Cl Co-Transporter Gene Slc12a2 Results in Deafness in Mice. Human Molecular Genetics 8:1579–1584. doi:10.1093/hmg/8.8.1579

Duvall III AJ. 1969. The ultrastructure of the external sulcus in the guinea pig cochlear duct. The Laryngoscope 79:1–29. doi:10.1288/00005537-196901000-00001

Forrest D, Erway LC, Ng L, Altschuler R, Curran T. 1996. Thyroid hormone receptor beta is essential for development of auditory function. Nat Genet 13:354–7. doi:10.1038/ng0796-354

Furness DN. 2019. Forgotten Fibrocytes: A Neglected, Supporting Cell Type of the Cochlea With the Potential to be an Alternative Therapeutic Target in Hearing Loss. Front Cell Neurosci 13:532. doi:10.3389/fncel.2019.00532

Galic M, Giebel W. 1989. An electron microscopic study of the function of the root cells in the external spiral sulcus of the cochlea. Acta Otolaryngol Suppl 461:1–15.

Grøntved L, Waterfall JJ, Kim DW, Baek S, Sung M-H, Zhao L, Park JW, Nielsen R, Walker RL, Zhu YJ, Meltzer PS, Hager GL, Cheng S. 2015. Transcriptional activation by the thyroid hormone receptor through ligand-dependent receptor recruitment and chromatin remodelling. Nat Commun 6:7048. doi:10.1038/ncomms8048

Groza T, Gomez FL, Mashhadi HH, Muñoz-Fuentes V, Gunes O, Wilson R, Cacheiro P, Frost A, Keskivali-Bond P, Vardal B, McCoy A, Cheng TK, Santos L, Wells S, Smedley D, Mallon A-M, Parkinson H. 2023. The International Mouse Phenotyping Consortium: comprehensive knockout phenotyping underpinning the study of human disease. Nucleic Acids Research 51:D1038–D1045. doi:10.1093/nar/gkac972

Gu S, Olszewski R, Taukulis I, Wei Z, Martin D, Morell RJ, Hoa M. 2020. Characterization of rare spindle and root cell transcriptional profiles in the stria vascularis of the adult mouse cochlea. Sci Rep 10:18100. doi:10.1038/s41598-020-75238-8

Hao X, Xing Y, Moore MW, Zhang J, Han D, Schulte BA, Dubno JR, Lang H. 2014. Sox10 expressing cells in the lateral wall of the aged mouse and human cochlea. PLoS One 9:e97389. doi:10.1371/journal.pone.0097389

Hebert JM, McConnell SK. 2000. Targeting of cre to the Foxg1 (BF-1) locus mediates loxP recombination in the telencephalon and other developing head structures. Dev Biol 222:296–306. doi:10.1006/dbio.2000.9732

Hibino H, Nin F, Tsuzuki C, Kurachi Y. 2010. How is the highly positive endocochlear potential formed? The specific architecture of the stria vascularis and the roles of the ion-transport apparatus. Pflugers Arch 459:521–33. doi:10.1007/s00424-009-0754-z

Jagger DJ, Forge A. 2015. Connexins and gap junctions in the inner ear--it’s not just about K(+) recycling. Cell Tissue Res 360:633–44. doi:10.1007/s00441-014-2029-z

Jagger DJ, Forge A. 2013. The enigmatic root cell - emerging roles contributing to fluid homeostasis within the cochlear outer sulcus. Hear Res 303:1–11. doi:10.1016/j.heares.2012.10.010

Jagger DJ, Forge A. 2006. Compartmentalized and signal-selective gap junctional coupling in the hearing cochlea. J Neurosci 26:1260–8. doi:10.1523/JNEUROSCI.4278-05.2006

Jagger DJ, Nevill G, Forge A. 2010. The Membrane Properties of Cochlear Root Cells are Consistent with Roles in Potassium Recirculation and Spatial Buffering. J Assoc Res Otolaryngol 11:435–48. doi:10.1007/s10162-010-0218-3

Jean P, Wong Jun Tai F, Singh-Estivalet A, Lelli A, Scandola C, Megharba S, Schmutz S, Roux S, Mechaussier S, Sudres M, Mouly E, Heritier A-V, Bonnet C, Mallet A, Novault S, Libri V, Petit C, Michalski N. 2023. Single-cell transcriptomic profiling of the mouse cochlea: An atlas for targeted therapies. Proceedings of the National Academy of Sciences 120:e2221744120. doi:10.1073/pnas.2221744120

Karolyi IJ, Dootz GA, Halsey K, Beyer L, Probst FJ, Johnson KR, Parlow AF, Raphael Y, Dolan DF, Camper SA. 2007. Dietary thyroid hormone replacement ameliorates hearing deficits in hypothyroid mice. Mamm Genome 18:596–608. doi:10.1007/s00335-007-9038-0

Kerr TP, Ross MD, Ernst SA. 1982. Cellular localization of Na+,K+-ATPase in the mammalian cochlear duct: significance for cochlear fluid balance. Am J Otolaryngol 3:332–338. doi:10.1016/s0196-0709(82)80006-9

Kim HJ, Gratton MA, Lee J-H, Perez Flores MC, Wang W, Doyle KJ, Beermann F, Crognale MA, Yamoah EN. 2013. Precise toxigenic ablation of intermediate cells abolishes the “battery” of the cochlear duct. J Neurosci 33:14601–14606. doi:10.1523/JNEUROSCI.2147-13.2013

Kimura RS. 1984. Sensory and accessory epithelia of the cochlea. Friedmann, I And J Ballantyne (Ed) Ultrastructural Atlas Of The Inner Ear Ix++329p Butterworths London, England; Boston, Mass, Usa Illus P101–132.

Lee JH, Chiba T, Marcus DC. 2001. P2X2 receptor mediates stimulation of parasensory cation absorption by cochlear outer sulcus cells and vestibular transitional cells. J Neurosci 21:9168–9174. doi:10.1523/JNEUROSCI.21-23-09168.2001

Li Y, Liu H, Zhao X, He DZ. 2020. Endolymphatic Potential Measured From Developing and Adult Mouse Inner Ear. Front Cell Neurosci 14:584928. doi:10.3389/fncel.2020.584928

Lim DJ, Anniko M. 1985. Developmental morphology of the mouse inner ear. A scanning electron microscopic observation. Acta Otolaryngol Suppl 422:1–69.

Liu S, Shen S, Yan Y, Sun C, Lu Z, Feng H, Ma Y, Tang Z, Yu J, Wu Y, Gereben B, Mohácsik P, Fekete C, Feng X, Yuan F, Guo F, Hu C, Shao M, Gao X, Zhao L, Li Y, Jiang J, Ying H. 2022. Triiodothyronine (T3) promotes brown fat hyperplasia via thyroid hormone receptor α mediated adipocyte progenitor cell proliferation. Nat Commun 13:3394. doi:10.1038/s41467-022-31154-1

Liu W, Wang C, Yu H, Liu S, Yang J. 2018. Expression of acetylated tubulin in the postnatal developing mouse cochlea. European journal of histochemistry: EJH 62. doi:10.4081/ejh.2018.2942

Marcus DC, Wu T, Wangemann P, Kofuji P. 2002. KCNJ10 (Kir4.1) potassium channel knockout abolishes endocochlear potential. Am J Physiol Cell Physiol 282:C403–407. doi:10.1152/ajpcell.00312.2001

Mendoza A, Tang C, Choi J, Acuña M, Logan M, Martin AG, Al-Sowaimel L, Desai BN, Tenen DE, Jacobs C, Lyubetskaya A, Fu Y, Liu H, Tsai L, Cohen DE, Forrest D, Wilson AA, Hollenberg AN. 2021. Thyroid hormone signaling promotes hepatic lipogenesis through the transcription factor ChREBP. Sci Signal 14:eabh3839. doi:10.1126/scisignal.abh3839

Mustapha M, Fang Q, Gong T-W, Dolan DF, Raphael Y, Camper SA, Duncan RK. 2009. Deafness and Permanently Reduced Potassium Channel Gene Expression and Function in Hypothyroid Pit1dw Mutants. The Journal of Neuroscience 29:1212. doi:10.1523/JNEUROSCI.4957-08.2009

Ng L, Cordas E, Wu X, Vella KR, Hollenberg AN, Forrest D. 2015. Age-Related Hearing Loss and Degeneration of Cochlear Hair Cells in Mice Lacking Thyroid Hormone Receptor beta1. Endocrinology 156:3853–65. doi:10.1210/en.2015-1468

Ng L, Kelley MW, Forrest D. 2013. Making sense with thyroid hormone--the role of T(3) in auditory development. Nat Rev Endocrinol 9:296–307. doi:10.1038/nrendo.2013.58

Roth B, Bruns V. 1992. Postnatal development of the rat organ of Corti. I. General morphology, basilar membrane, tectorial membrane and border cells. Anat Embryol (Berl) 185:559–69. doi:10.1007/BF00185615

Royaux IE, Belyantseva IA, Wu T, Kachar B, Everett LA, Marcus DC, Green ED. 2003. Localization and functional studies of pendrin in the mouse inner ear provide insight about the etiology of deafness in pendred syndrome. J Assoc Res Otolaryngol 4:394–404. doi:10.1007/s10162-002-3052-4

Rüsch A, Erway LC, Oliver D, Vennström B, Forrest D. 1998. Thyroid hormone receptor β-dependent expression of a potassium conductance in inner hair cells at the onset of hearing. Proceedings of the National Academy of Sciences 95:15758–15762. doi:10.1073/pnas.95.26.15758

Rusch A, Ng L, Goodyear R, Oliver D, Lisoukov I, Vennstrom B, Richardson G, Kelley MW, Forrest D. 2001. Retardation of cochlear maturation and impaired hair cell function caused by deletion of all known thyroid hormone receptors. J Neurosci 21:9792–9800. doi:10.1523/JNEUROSCI.21-24-09792.2001

Sakagami M, Fukazawa K, Matsunaga T, Fujita H, Mori N, Takumi T, Ohkubo H, Nakanishi S. 1991. Cellular localization of rat Isk protein in the stria vascularis by immunohistochemical observation. Hear Res 56:168–172. doi:10.1016/0378-5955(91)90166-7

Sawant O, Horton AM, Shukla M, Rayborn ME, Peachey NS, Hollyfield JG, Rao S. 2015. Light-Regulated Thyroid Hormone Signaling Is Required for Rod Photoreceptor Development in the Mouse Retina. Invest Ophthalmol Vis Sci 56:8248–8257. doi:10.1167/iovs.15-17743

Sharlin DS, Ng L, Verrey F, Visser TJ, Liu Y, Olszewski RT, Hoa M, Heuer H, Forrest D. 2018. Deafness and loss of cochlear hair cells in the absence of thyroid hormone transporters Slc16a2 (Mct8) and Slc16a10 (Mct10). Sci Rep 8:4403. doi:10.1038/s41598-018-22553-w

Shodo R, Hayatsu M, Koga D, Horii A, Ushiki T. 2017. Three-dimensional reconstruction of root cells and interdental cells in the rat inner ear by serial section scanning electron microscopy. Biomed Res 38:239–248. doi:10.2220/biomedres.38.239

Song MH, Choi S-Y, Wu L, Oh S-K, Lee HK, Lee DJ, Shim D-B, Choi JY, Kim U-K, Bok J. 2011. Pou3f4 deficiency causes defects in otic fibrocytes and stria vascularis by different mechanisms. Biochemical and Biophysical Research Communications 404:528–533. doi:10.1016/j.bbrc.2010.12.019

Soriano P. 1999. Generalized lacZ expression with the ROSA26 Cre reporter strain. Nat Genet 21:70–71. doi:10.1038/5007

Spicer SS, Schulte BA. 1996. The fine structure of spiral ligament cells relates to ion return to the stria and varies with place-frequency. Hear Res 100:80–100. doi:10.1016/0378-5955(96)00106-2

Steel KP, Barkway C. 1989. Another role for melanocytes: their importance for normal stria vascularis development in the mammalian inner ear. Development 107:453–463. doi:10.1242/dev.107.3.453

Sun J, Ahmad S, Chen S, Tang W, Zhang Y, Chen P, Lin X. 2005. Cochlear gap junctions coassembled from Cx26 and 30 show faster intercellular Ca2+ signaling than homomeric counterparts. American Journal of Physiology-Cell Physiology 288:C613–C623. doi:10.1152/ajpcell.00341.2004

Umesono K, Murakami KK, Thompson CC, Evans RM. 1991. Direct repeats as selective response elements for the thyroid hormone, retinoic acid, and vitamin D3 receptors. Cell 65:1255–1266. doi:10.1016/0092-8674(91)90020-Y

Uziel A. 1986. Periods of sensitivity to thyroid hormone during the development of the organ of Corti. Acta Otolaryngol Suppl 429:23–27. doi:10.3109/00016488609122726

Uziel A, Legrand C, Ohresser M, Marot M. 1983. Maturational and degenerative processes in the organ of Corti after neonatal hypothyroidism. Hear Res 11:203–218. doi:10.1016/0378-5955(83)90079-5

Uziel Alain, Legrand C, Rabie A. 1985. Corrective effects of thyroxine on cochlear abnormalities induced by congenital hypothyroidism in the rat. I. Morphological study. Developmental Brain Research 19:111–122. doi:10.1016/0165-3806(85)90236-6

Walters BJ, Zuo J. 2013. Postnatal development, maturation and aging in the mouse cochlea and their effects on hair cell regeneration. Hear Res 297:68–83. doi:10.1016/j.heares.2012.11.009

Wangemann P. 2006. Supporting sensory transduction: cochlear fluid homeostasis and the endocochlear potential. J Physiol 576:11–21. doi:10.1113/jphysiol.2006.112888

Winter H, Rüttiger L, Müller M, Kuhn S, Brandt N, Zimmermann U, Hirt B, Bress A, Sausbier M, Conscience A, Flamant F, Tian Y, Zuo J, Pfister M, Ruth P, Löwenheim H, Samarut J, Engel J, Knipper M. 2009. Deafness in TRbeta mutants is caused by malformation of the tectorial membrane. J Neurosci 29:2581–2587. doi:10.1523/JNEUROSCI.3557-08.2009

Xie Z, Ma X, Ji W, Zhou G, Lu Y, Xiang Z, Wang YX, Zhang L, Hu Y, Ding Y-Q, Zhang WJ. 2010. Zbtb20 is essential for the specification of CA1 field identity in the developing hippocampus. Proc Natl Acad Sci U S A 107:6510–6515. doi:10.1073/pnas.0912315107

Xie Z, Ma X-H, Bai Q-F, Tang J, Sun J-H, Jiang F, Guo W, Wang C-M, Yang R, Wen Y-C, Wang F-Y, Chen Y-X, Zhang H, He DZ, Kelley MW, Yang S, Zhang WJ. 2023. ZBTB20 is essential for cochlear maturation and hearing in mice. Proceedings of the National Academy of Sciences 120:e2220867120. doi:10.1073/pnas.2220867120

Xie Z, Zhang H, Tsai W, Zhang Y, Du Y, Zhong J, Szpirer C, Zhu M, Cao X, Barton MC, Grusby MJ, Zhang WJ. 2008. Zinc finger protein ZBTB20 is a key repressor of alpha-fetoprotein gene transcription in liver. Proc Natl Acad Sci U S A 105:10859–10864. doi:10.1073/pnas.0800647105

Yan Y, Niu Z, Sun C, Li P, Shen S, Liu S, Wu Y, Yun C, Jiao T, Jia S, Li Yuying, Fang Z-Z, Zhao L, Wang J, Xie C, Jiang C, Li Yan, Feng X, Hu C, Jiang J, Ying H. 2022. Hepatic thyroid hormone signalling modulates glucose homeostasis through the regulation of GLP-1 production via bile acid-mediated FXR antagonism. Nat Commun 13:6408. doi:10.1038/s41467-022-34258-w

Zilz ND, Murray MB, Towle HC. 1990. Identification of multiple thyroid hormone response elements located far upstream from the rat S14 promoter. J Biol Chem 265:8136–8143.

